# Metabolic adaptations underlie epigenetic vulnerabilities in chemoresistant breast cancer

**DOI:** 10.1101/286054

**Authors:** Geneviève Deblois, Seyed Ali Madani Tonekaboni, Yunchi Ingrid Kao, Felicia Tai, Xiaojing Liu, Ilias Ettayebi, Paul Guilhamon, Alexandra Fedor, Giacomo Grillo, Alexander Murison, Wail Ba alawi, David Cescon, Cheryl Arrowsmith, Daniel DeCarvalho, Morag Park, Benjamin Haibe-Kains, Jason W. Locasale, Mathieu Lupien

**Affiliations:** Princess Margaret Cancer Centre, University Health Network, Toronto, ON, M5G 1L7, Canada; Department of Medical Biophysics, University of Toronto, Toronto, ON, M5G 1L7, Canada; Department of Computer Sciences, University of Toronto, Toronto, ON, Canada; Department of Pharmacology and Cancer Biology, Duke University School of Medicine, Durham, NC 27710, USA; Structural Genomics Consortium, Toronto, ON M5G 1L7, Canada; Goodman Cancer Research Centre, McGill University, Montreal, QC, H3A 1A3, Canada; Ontario Institute for Cancer Research, Toronto, ON M5G 0A3, Canada

**Keywords:** Drug resistance, paclitaxel, methionine, S-adenosylmethionine, epigenetic inhibitor, DNA methylation, H3K27me3, dsRNA, chromatin, epigenetic, breast cancer, transposable elements, endogenous retroviral elements, transposon, EZH2 inhibitor, histone modification, triple-negative breast cancer

## Abstract

Cancer cell survival upon cytotoxic drug exposure leads to changes in cell identity, dictated by the epigenome. Several metabolites serve as substrates or co-factors to chromatin-modifying enzymes, suggesting that metabolic changes can underlie change in cell fate. Here, we show that progression of triple-negative breast cancer (TNBC) to taxane-resistance is characterized by altered methionine metabolism and S-adenosylmethionine (SAM) availability, giving rise to DNA hypomethylation in regions enriched for transposable elements (TE). Compensatory redistribution of H3K27me3 forming Large Organized Chromatin domains of lysine (K) modification (LOCK) prevents expression of TE in taxane-resistant cells. Pharmacological inhibition of EZH2, the H3K27me3 methyltransferase, alleviates TE repression, leading to the accumulation of dsRNA and activation of the interferon viral mimicry-response, specifically inhibiting the growth of taxane-resistant TNBC. Together, our work delineates a role for metabolic adaptations in redefining the epigenome of taxane-resistant TNBC cells and underlies an epigenetic vulnerability toward pharmacological inhibition of EZH2.

## INTRODUCTION

Despite initial benefits of cytotoxic chemotherapy such as taxanes for the treatment of triple-negative breast cancer patients (TNBC), clinical drug resistance is a major barrier to remission(Broxterman et al., 2009). Intrinsic or acquired therapeutic resistance in TNBC results in drug-tolerant cell populations with the capacity to survive to the toxic environment that stems from exposure to drugs(Ramirez et al., 2016; Sharma et al., 2010). Targeting the initial events that contribute to the adaptation of cancer cells to such environments, including metabolic and epigenetic reprogramming, may help prevent the occurrence of drug-resistant cell populations.

Epigenetic modifications such as DNA methylation and histone modifications directly regulate cell identity and genomic stability by controlling the accessibility of chromatin to the transcriptional machinery(Adam and Fuchs, 2016). Reprogramming of epigenetic modifications has extensively been implicated as drivers of carcinogenesis(Mack et al., 2014; Momparler and Bovenzi, 2000; Sharma et al., 2010; Varier and Timmers, 2011; Yao et al., 2015) and also typify cancer progression to therapeutic resistance. This is exemplified by the changes in histone 3 (H3) lysine 4 dimethylation (H3K4me2) and in H3 lysine 36 trimethylation (H3K36me3) distribution in luminal breast cancer cells progressing from endocrine-therapy sensitivity to resistance(Magnani et al., 2013) and by the evolution of distant metastasis in pancreatic cancer characterized by methylation of H3 lysine 9 forming Large Organized Chromatin H3 lysine (K) 9 (H3K9) modification domains (LOCKs)(McDonald et al., 2017). This suggests that the reprogramming of epigenetic landscapes can contribute a selective advantage for cancer cells to better survive exposure to cytotoxic drugs.

Metabolic adaptation underlies energetic, biomass and detoxification requirements for cancer cell survival(Pavlova and Thompson, 2016). In addition, altered glucose metabolism, amino-acid dependency or increased cellular antioxidant capacity are metabolic adaptations enabling cancer cell survival upon exposure to cytotoxic drugs and favoring the development of drug-resistant cancer cell populations(Ju et al., 2015; Kerr et al., 2016; Komurov et al., 2012). Metabolism can directly impact epigenetic processes since specific metabolites serve as substrates or co-factors for epigenetic modifications(Lu and Thompson, 2012; Reid et al., 2017). In particular, methylation of DNA and histones relies on the availability of the universal methyl donor metabolite S-adenosylmethionine (SAM), which is biosynthesized from the precursor L-methionine as part of the one-carbon metabolism(Mentch et al., 2015). Accordingly, disruption of the one-carbon metabolism through the induction of the serine-glycine pathway in LKB-1-deficient pancreatic tumors raises SAM level with concomitant increase in DNA methylation(Kottakis et al., 2016). Furthemore, SAM-consuming metabolic pathways including biosynthesis of polyamines, phosphatidylcholine, phospholipids, carnitine and 1-MNA that are prevalent in cancer cells can sequester SAM away from histone and DNA methylation(Martínez-Uña et al., 2013; Obeid and Herrmann, 2009; Ulanovskaya et al., 2013; Venkatachalam, 2015; Ye et al., 2017). Because the one-carbon metabolism is the sensor of nutritional and environmental status(Locasale, 2013; Newman and Maddocks, 2017), its disruption can account for epigenetic alterations that allow cancer cells adaptations to environmental changes such as toxic exposure to drugs.

Here we report that taxane-resistant TNBC cells undergo metabolic adaptations affecting one-carbon metabolism and SAM availability, resulting in DNA hypomethylation and concomitant compensatory reprogramming of H3K27me3 distribution across the genome into large chromatin domains reminiscent of LOCKs. This epigenetic reprogramming provides a compensatory mechanism ensuring repression of TE at H3K27me3 LOCKs. Finally, we demonstrate that the switch from DNA methylation to H3K27me3-mediated repression over transposable elements creates a vulnerability of taxane-resistant TNBC cells to epigenetic therapy based on EZH2 inhibitors, allowing for double stranded RNA production and activation of the viral mimicry response.

## RESULTS

### One-carbon metabolism is altered in taxane-resistant TNBC cells

To assess the contribution of metabolic adaptations to taxane resistance in TNBC we derived six independent paclitaxel-resistant models from MDA-MB-436 (resistant population MDA-MB-436-R20A, R20B and R20C) and Hs 578T TNBC cell lines (resistant population Hs 578T-R20A, R20B and R10C) upon long-term exposure to increasing concentrations of paclitaxel (Figure S1A). Dose and time-dependent cell viability assays show a 10 to 30-fold reduced sensitivity toward paclitaxel or docetaxel, another taxane derivative, in resistant versus sensitive cells (Figure 1A, B) and clear discrimination in the sensitivity at the 20 nM working paclitaxel concentration for the resistant populations relative to parental cells (Figure S1B and S1C). Dose-dependent cell viability assays using non-taxane drugs including 5-FU, doxorubicin and cisplatin reveal no cross-resistance in the taxane-resistant TNBC populations (Figure 1B and S1D), suggesting that the six taxane-resistant TNBC populations rely on a taxane-specific resistance mechanism as opposed to a generic mechanism of drug resistance. Gene expression profiling using RNA-seq for the 6 taxane-resistant populations and parental cells reveals significant downregulation in the expression of genes involved in interferon signaling, immune response, adhesion, differentiation/development and cell communication in the taxane-resistant populations relative to their respective parental cells, suggesting that the resistance is unlikely due to drug-related pharmacological changes in our models (Figure S1E, S1F and S1G).

**Figure 1.**
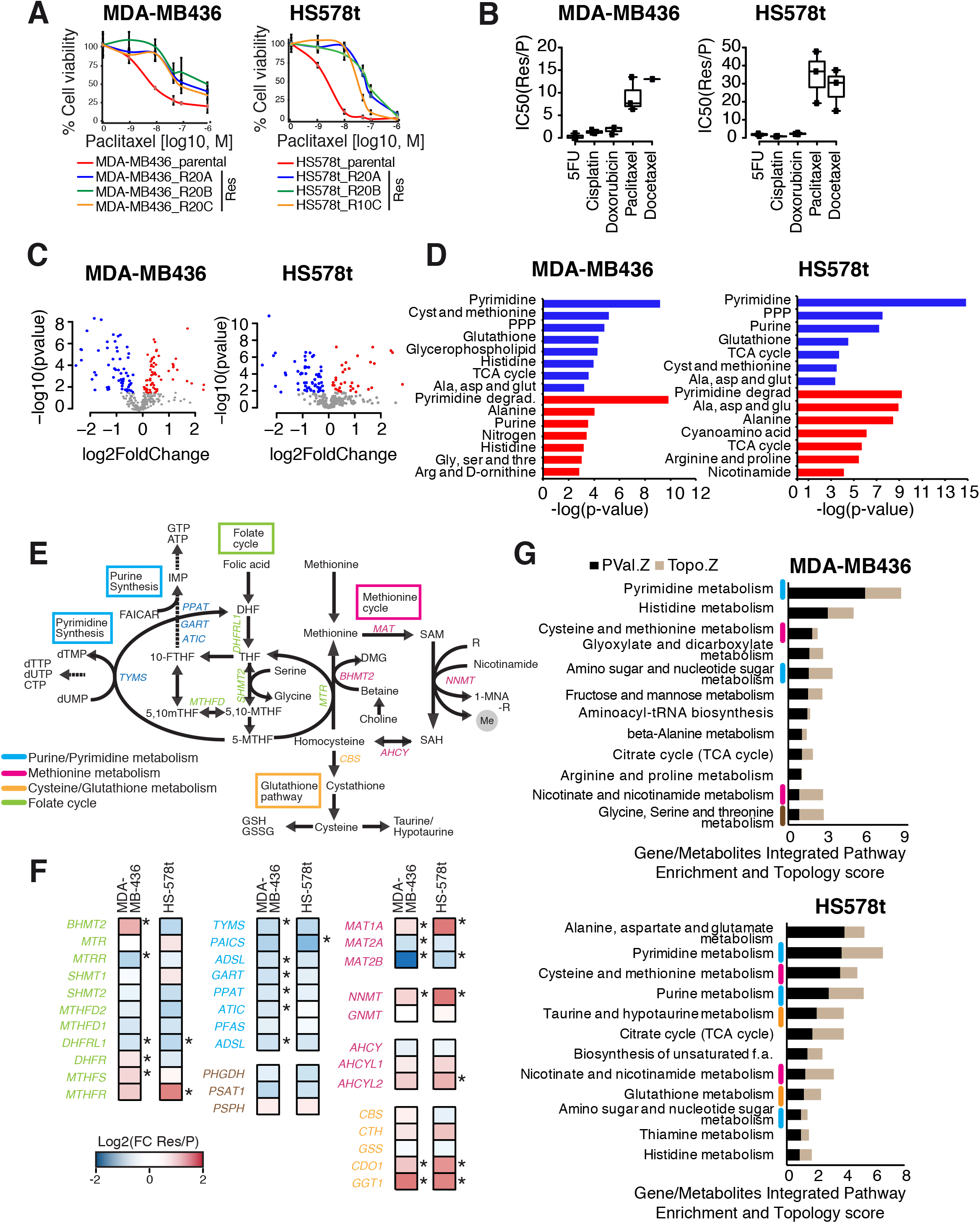
The one-carbon metabolism is altered in taxane-resistant TNBC cells. (A) Forty-eight hours paclitaxel dose-response curves of taxane-resistant derived cell populations and of parental MDA-MB-436 and Hs 578T cells. Error bars: standard deviation. (B) Boxplot showing the relative IC50 of resistant cells to IC50 of parental cells upon 48h treatment with the standard-of-care chemotherapeutic agents 5’FU, Cisplatin, Doxorubicin, Docetaxel and Paclitaxel. (C) Volcano plots depicting fold change ratio of metabolites significantly altered in paclitaxel-resistant cells relative to parental cells. Each dot represents a metabolite (p<0.05, statistical significance calculated by one-way analysis of variance (ANOVA); metabolites in gray are not significant, metabolites in blue are significantly decreased in taxane-resistant cells, metabolites in red are significantly increased in taxane-resistant cells). (D) Histogram of metabolite pathway enrichment using human Kyoto Encyclopedia of Genes and Genome (KEGG) compound database and Small Molecule Pathway Database (SMPDB) in taxane-resistant TNBC cells relative to sensitive cells. Blue bars represent pathways significantly enriched among donwregulated metabolites, bars represent pathways significantly enriched among upregulated metabolites (E) Schematic representation of the one-carbon unit cycle and the various metabolic pathways that it comprises. (F) Relative gene expression from RNA-seq data for taxane-resistant cells relative to parental cells for genes involved in the metabolic reactions of the one-carbon unit cycle. Statistical significance calculated using the BH-adjusted FDR. *, FDR<0.05). (G) Integrated gene-metabolites pathway enrichment using human KEGG pathway enrichment for metabolites and gene expression. Black bars represent Z-score for pathway enrichment in integrated genes-metabolites data (Hypergeometric test); beige bar represent Z-score for topology analysis of gene-metabolites (Degree centrality).

We next delineated the metabolic adaptations that characterize the acquisition of taxane-resistance in TNBC using metabolomics profiling with liquid chromatography-high resolution mass spectrometry (LC-HRMS) of intracellular metabolite extracts from all six taxane-resistant and both parental TNBC cells. This identified 69 and 62 metabolites whose levels were significantly decreased in taxane-resistant MDA-MB-436 and Hs 578T cells respectively while 67 and 29 metabolites showed significantly increased levels in taxane-resistant MDA-MB-436 and Hs 578T cells respectively, relative to their corresponding parental cells (Figure 1C). Metabolic pathway enrichment analysis using the human Kyoto Encyclopedia of Genes and Genome (KEGG) compound database and Small Molecule Pathway Database (SMPDB) revealed significant depletion of metabolites involved in purine/pyrimidine, glutathione and in cysteine and methionine metabolism in taxane-resistant MDA-MB-436 and Hs 578T TNBC cells compared to parental cells (Figure 1D and Figure S1H). On the other hand, we observed an increase in the abundance of a few metabolites in the purine/pyrimidine pathway and alanine metabolism across all taxane-resistant models relative to parental cells (Figure 1D and Figure S1H).

These altered metabolic pathways are interconnected through the one-carbon metabolic pathway (Figure 1E). Gene expression profiling in taxane-resistant TNBC cells relative to parental cells using RNA-seq data further highlights the dysregulation of the one-carbon metabolism as illustrated by the differential expression of key genes regulating metabolic enzymes controlling methionine, and purine/pyrimidine metabolism in taxane-resistant cells relative to parental cells, including *MAT2A*, *MAT2B*, *TYMS* and *ADSL* genes (Figure 1F). Integration of metabolomic and gene expression profiles from the taxane-resistant TNBC cells relative to parental cells demonstrate the significant enrichment of the one-carbon-related metabolic pathways (Figure 1G). Overall, our results suggest that the development of taxane-resistance in our TNBC cells converge on metabolic adaptations affecting various aspects of the one-carbon metabolism, inclusive of the methionine cycle.

### Taxane-resistant TNBC cells display impaired methionine metabolism

Through the recycling of methionine, the one-carbon metabolism contributes to the biosynthesis of key metabolites for cancer cells, including that of SAM, the universal methyl group donor required for DNA and histone methylation (Figure 2A). The LC-HRMS results reveal a significant decrease (p-value < 0.05) in the steady-state levels of most metabolites part of the methionine metabolism in taxane-resistant cells relative to parental cells, including SAM, SAH, L-homocysteine and L-methionine (Figure 2B). Conversion ratios across successive metabolites in the methionine cycle, indicative of changes in metabolic flux, reveal a significant decrease in the ratio of L-homocysteine (Hcy) to L-methionine (Met) (Hcy/Met) in all models of taxane-resistance relative to their respective parental TNBC cells (Figure 2C) while the ratios between other successive metabolites are not uniformly affected across the resistant cells (Figure S2A). This is suggestive of alterations in the activity of methionine cycle and in the usage of methionine-related metabolites in taxane-resistant TNBC cells relative to parental cells. Accordingly, the conversion ratio between Hcy and cysteine (Cyst/Hcy) is significantly elevated in the taxane-resistant cells, indicative of increased usage of Hcy towards the transufuration/glutathione pathway, likely contributing to metabolic detoxification reactions (Figure 2D). This is further supported by a significant restoration of the detoxification potential defined by the ratios of reduced over oxidized glutathione (GSH/GSSG) in the taxane-resistant TNBC cells relative to taxane-treated parental cells (Figure S2B).

**Figure 2.**
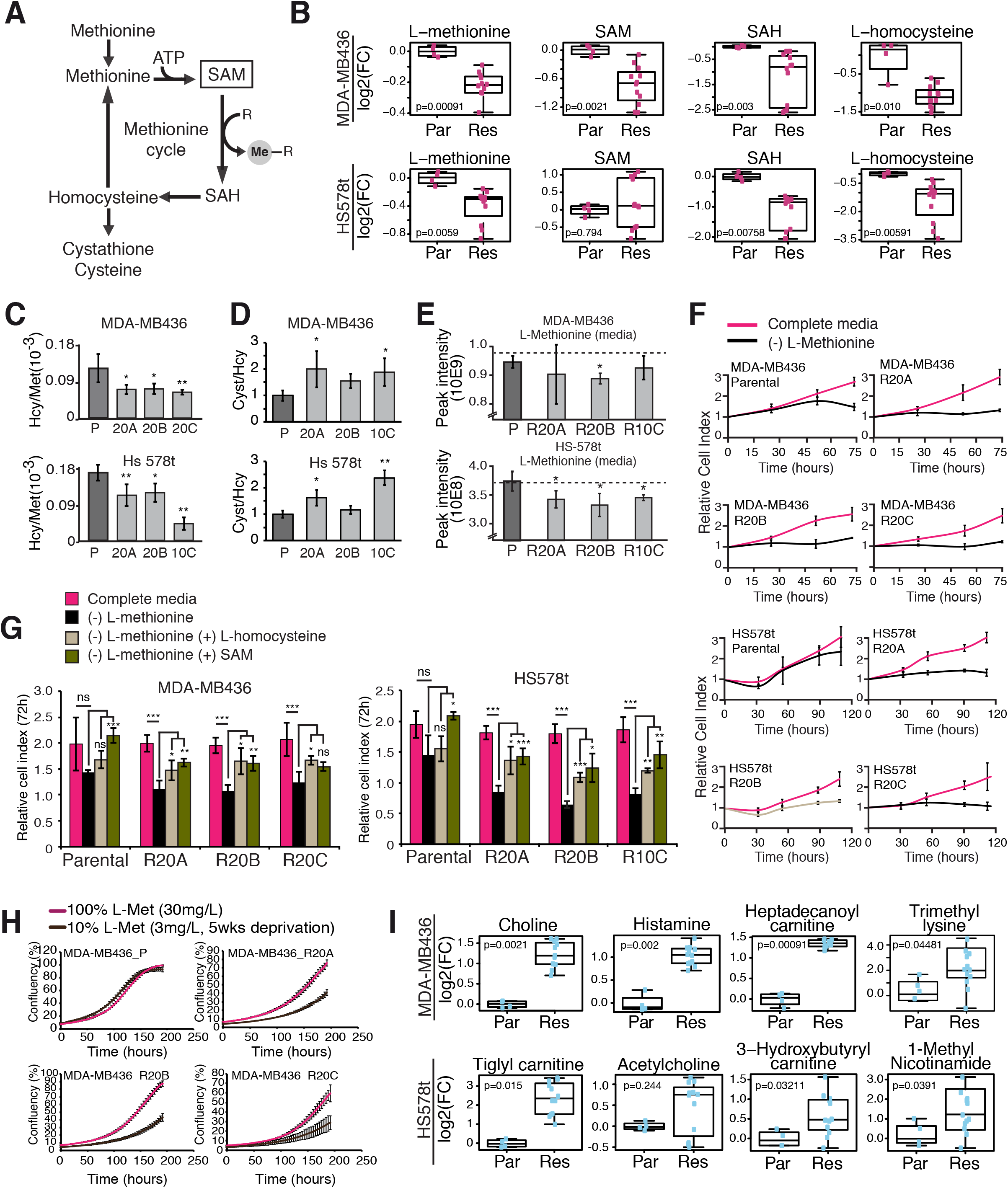
Taxane-resistant TNBC cells display impaired methionine metabolism. (A) Schematic representation of the methionine cycle coupled to the folate cycle. (B) Box plot of fold-change in steady-state intracellular methionine cycle metabolites in taxane-resistant relative to parental cells. Statistical significance calculated using one-way ANOVA (p<0.05). (C) Ratio of relative abundance in steady-state levels of L-homocysteine (L-Hcy) to L-Methionine (L-Met) in parental and taxane-resistant TNBC cells. Error bars show standard deviation (std), n=4, statistical significance is calculated using 2-sided unpaired t-test, * denotes p<0.05; ** denotes p<0.01. (D) Ratio of relative abundance in steady-state levels of L-Hcy to Cysteine (Cyst) in parental and taxane-resistant TNBC cells. Error bars show standard deviation (std), n=4, statistical significance is calculated using 2-sided unpaired t-test, * denotes p<0.05; ** denotes p<0.01. (E) Histogram showing LC-MS peak intensity signal representative of the relative steady-state abundance of L-methionine in the conditioned media (3 days) in both parental and resistant cells. Dashed line represents the metabolite levels in naïve media. Error bars show std deviation, statistical significance is calculated using 2-sided unpaired t-test, * denotes p<0.05. (F) Time-dependent growth curves showing relative cell index over 75 hours for taxane-resistant and parental MDA-MB-436 and Hs 578T cells upon complete depletion of L-methionine from the growth media. Error bars: std; * denotes p<0.05; ** denotes p<0.01 by two-sided unpaired t-test. G) Relative cell index at a single time point (48h) depicting confluency at 48 hours upon methionine depletion and add-back of either L-methionine (30 mg/L), L-homocysteine (0.4 mM) or SAM (250 μM) in the growth media. Error bars: std deviations. Statistical significance was calculated by two-sided unpaired t-test, * denotes p<0.05; ** denotes p<0.01. (H) Time-dependent growth assay (IncucyteZoom) of parental and taxane-resistant MDA-MB-436 cells cells upon long term (5 weeks) deprivation of L-methionine (using 3 mg/L, 10% of initial amount). Resistant cells were exposed to paclitaxel (20 nM) at all time. Error bars: std. (I) Box plot showing fold-change difference in steady-state levels of SAM-consuming metabolites in taxane-resistant relative to parental cells. Statistical significance calculated using one-way ANOVA (p<0.05).

We next assessed whether taxane-resistant cells compensate for this defect in methionine recycling by increasing methionine uptake from the media. In agreement, LC-HRMS measurements using 48 hours conditioned media compared to naive media reveal a significant decrease in the total levels of L-methionine in most taxane-resistant TNBC cells relative to their respective parental cells (Figure 2E P<0.05, 2-sided unpaired t-test). This led us to test the dependence of taxane-resistant cells on L-methionine for proliferation. Cell proliferation assay revealed that the growth of taxane-resistant TNBC cells was significantly impaired upon complete depletion of L-methionine from the culture media (Figure 2F, P<0.05 as of 48 hours) while only a moderate decrease in relative growth of parental cells was observed at 48 hours following L-methionine depletion (Figure 2F). Supplementation with SAM and L-homocysteine partially rescued the growth defect in taxane-resistant TNBC cells deprived of L-methionine (Figure 2G), suggesting that the increased requirement for L-methionine uptake contributes to the recycling of methionine in order to sustain SAM and Hcy production in taxane-resistant TNBC cells. Likewise, a significant decrease (P<0.001 at end point) in the growth of taxane-resistant MDA-MB-436 cells relative to parental cells upon long term (5 weeks) culturing in growth media partially-deprived of L-methionine (containing 3 μg/L L-methionine, 10% of normal levels) (Figure 2H) further support increased dependency towards methionine uptake for proper methionine recycling in the taxane-resistant cells.

SAM contributes to various biological reactions in cells (Figure S2C). In order to identify the possible fates of SAM in the taxane-resistant cells that display altered methionine recycling, we assessed the conversion ratios for successive metabolites involved in SAM-consuming metabolic pathways including synthesis of carnitine, 1-methylnicotinamide (1-MNA) and choline. The ratios of carnitine:lysine and 1-MNA:NAD are significantly increased in Hs 578T cells (Figure S2D-F). Likewise, an increase in the choline-derivative:ethanolamine-derivative ratios is observed in the taxane-resistant MDA-MB-436 cells relative to parental cells (Figure S2E). These observations indicate increased usage of SAM in SAM-consuming metabolic pathways in taxane-resistant cells relative to parental cells. Accordingly, metabolomic profiling revealed an increase in the steady-state intracellular levels of choline, choline derivatives and carnitine-derivatives in MDA-MB-436 taxane-resistant cells and in carnitine-derivatives and 1-methylnicotinamide in taxane-resistant Hs 578T cells relative to the parental TNBC cells (P <0.05, Figure 2I). Overall, metabolomic analyses show that altered one-carbon metabolism in taxane-resistant in TNBC cells is characterized impaired methionine recycling that stem from increased usage of SAM in SAM-consuming metabolic pathways including transulfuration and synthesis of 1-MNA, choline and carnitine and resulting in increased dependency on exogenous methionine.

### Altered SAM levels lead to DNA hypomethylation at intergenic regions in taxane-resistant TNBC cells

Due to the observed increase in SAM-consuming metabolites, coupled with the global decrease in steady-state levels of metabolites involved in methionine metabolism, we assessed the levels of DNA and histone methylation in taxane-resistant and parental TNBC cells. Using an enzyme-linked immunosorbant assay (ELISA) system that assesses global DNA methylation using long interspersed nuclear element (LINE-1) probe, we observed a significant decrease in the levels of global DNA methylation in taxane-resistant MDA-MB-436 and Hs 578T cells relative to parental cells (Figure 3A). We next tested for specific changes in DNA methylation in taxane-resistant and parental MDA-MB-436 cells using the EPIC methylarray comprising 850,000 CpG probes. We observed a change in the distribution of DNA methylation signal across the genome of taxane-resistant compared to parental MDA-MB-436 cells illustrated by decreased Pearson correlation score (Figure 3B). In particular, 29,247 significantly hypomethylated versus 12,283 significantly hypermethylated CpGs were identified in taxane-resistant relative to parental TNBC cells (delta-beta-value cutoff +/- 0.3 and *P*<0.05) (Figure 3C). Identification of differentially methylated regions (DMRs) using Probe Lasso method reveals similar results with 1559 significantly hypomethylated DMRs versus 374 hypermethylated DMRs in taxane-resistant versus parental TNBC cells (Figure S3A).

**Figure 3.**
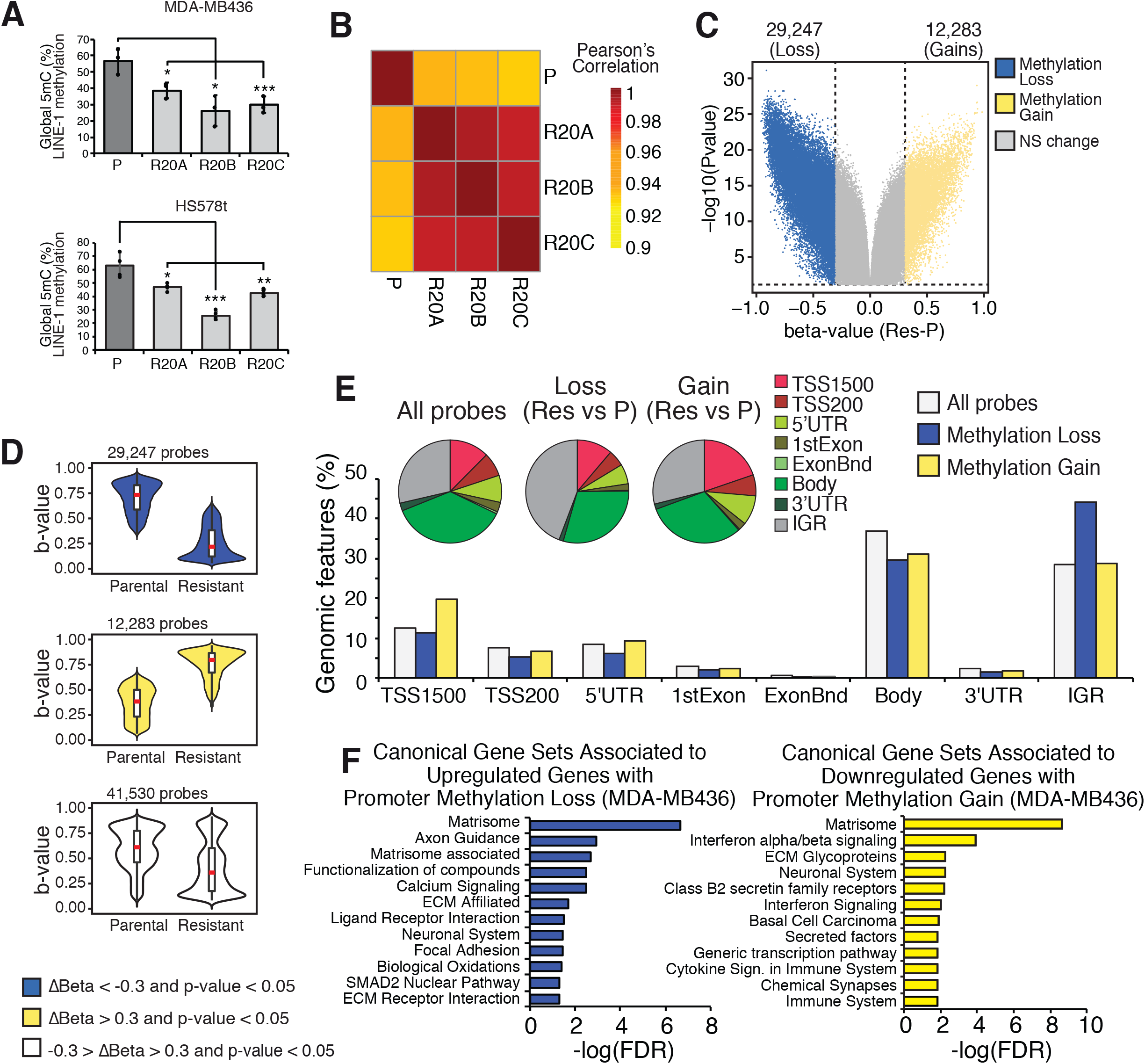
Altered SAM levels lead to DNA hypomethylation at intergenic regions in taxane-resistant TNBC cells. (A) Global 5mC levels (% of total cytosine) at the steady state level in both parental and taxane-resistant cells as assayed by Global 5mC methylation ELISA assay on LINE-1 probes. Error bars: std, statistical significance calculated by two-sided unpaired t-test, * denotes p<0.05; *** denotes p<0.001. (B) Heat map of Pearson correlation score of DNA methylation beta-value distribution in parental and resistant MDA-MB-436 cells normalized 5mC beta-value as assessed by bisulfite conversion and hybridization to Methyl-EPIC 850K array and analyzed using Bioconductor ChAMP package. (C) Volcano plot showing significant differentially methylated probes in taxane-resistant relative to parental MDA-MB-436 cells (delta-beta-value (Resv-P)); (p<0.05). Each dot represents a differentially methylated probe, blue dots represent significantly hypomethylated CpGs with delta-beta value < −0.3 (Res-P), yellow dots represent significantly hypomethylated CpGs with delta-beta value > +0.3 (Res-P), gray dots represent non-significantly (NS) altered CpGs with delta-beta values between −0.3 and +0.3. (D) Violin plots representing the distribution of delta-beta values (Res-P) in parental and taxane-resistant MDA-MB-436 cells for either all significantly altered CpGs (bottom panel) or for significantly hypomethylated CpGs (top panel) or for hypermethylated CpGs (middle panel) (E) Percentage of distribution of genomic features associated to all probes (white), to the 29247 probes showing a significant loss of methylation in the resistant cells (blue) or to the 12283 probes showing a significant gain of methylation in resistant cells relative to parental cells. Insert represents venn diagram of the distribution of the genomic features in each subtypes. (F) Canonical gene-set enrichment associated to upregulated genes that have promoter 5mC loss (blue) or with downregulated genes that have promoter 5mC gain in the resistant cells relative to parental cells.

The gains and losses in DNA methylation events considered are representative of changes between low methylation and high methylation states and dismiss small variations that could originate from technical issues (Figure 3D top 2 panels and Figure S3B). The significantly higher number of hypomethylated CpGs relative to hypermethylated CpGs reveals an overall global decrease in DNA methylation in taxane-resistant MDA-MB-436 cells relative to parental cells (Figure 3D bottom panel). The hypomethylated CpGs in taxane-resistant cells are enriched at intergenic (IGR) regions relative to the distribution of all CpG assessed, whereas hypermethylated CpGs in taxane-resistant cells are mostly observed at promoter regions (TSS1500) (Figure 3E and S3C).

To infer the impact of the reprogramming of DNA methylation on gene expression, we assessed enrichment of gene ontology pathways for differentially regulated genes with hypomethylated or hypermethylated promoters (TSS1500, TSS200, 5’UTR) in taxane-resistant MDA-MB-436 cells. Downregulated genes harbouring hypermethylated promoters in taxane-resistant TNBC cells were significantly associated with enrichment of the interferon alpha/beta signaling, immune system and cytokine signaling gene sets (Figure 3F and S3D-E). Overall, these results demonstrate a reprogramming of DNA methylation operating in taxane-resistant TNBC and resulting in a significant depletion of DNA methylation over intergenic regions.

### The distribution of H3K27me3 is reprogrammed in taxane-resistant TNBC cells

To assess if changes in SAM levels have an impact on the chromatin beyond DNA methylation, we assessed the global levels of key histone methylations, namely the H3K4me1, H3K4me3, H3K27me1 and H3K27me3, by immunoblotting of total histone acid extracts, H3K27ac and global H3 serving as controls. The global levels of the various histone methylation marks were mostly unaffected in the taxane-resistant MDA-MB-436 cells relative to the parental sensitive cells (Figure 4A and S4A). Similar results were observed for histones purified from total cell extract using RIPA lysis buffer and sonication (Figure S4B). However, the H3K27me3 modification was markedly decreased in taxane-resistant relative to parental TNBC cells when using a mild modified RIPA lysis buffer that excludes the insoluble proteins presumably linked to heterochromatin(Henikoff et al., 2009) (Figure 4B and S4C). No difference was observed under these conditions for the H3K27ac modification (Figure S4A-B-C). ELISA-quantification of H3K27me3 modification from total histone acid extraction (Figure S4D) confirmed no significant change in the global levels of H3K27me3 when assaying total histone extracts. Overall, these observations suggest that changes in SAM levels in taxane-resistant TNBC cells does not lead to significant change in the histone lysine methylation assessed but is associated to a disruption of H3K27me3 levels away from soluble towards insoluble fractions, indicative of H3K27mne3 reprogramming.

**Figure 4.**
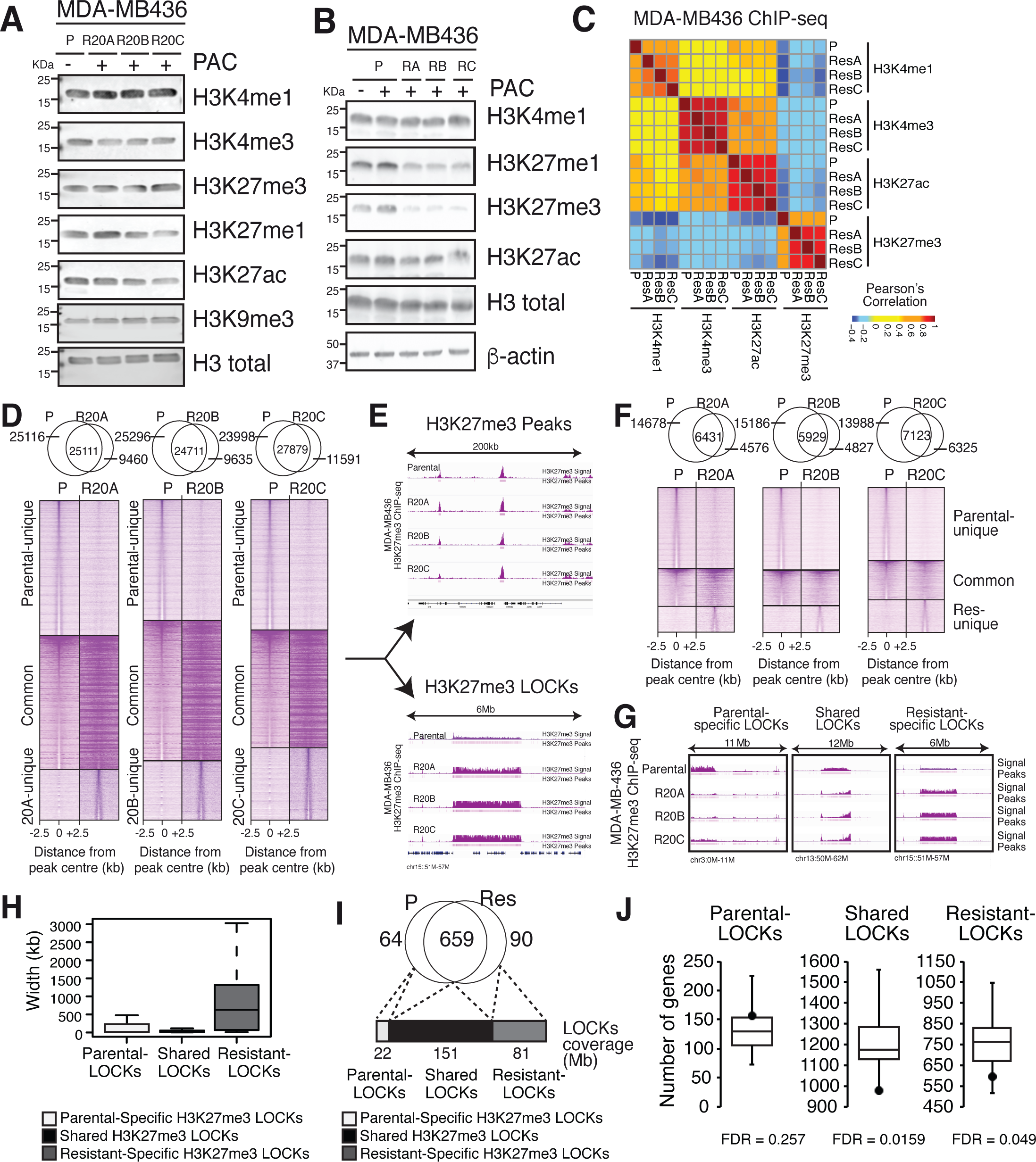
The distribution of H3K27me3 is reprogrammed in taxane-resistant TNBC cells. (A) Western blot depicting the levels of histone modifications as indicated in taxane-resistant and sensitive MDA-MB-436 cells. Cell extract was obtained using acidic extraction of histones. PAC, paclitaxel (20 nM). (B) Western blot depicting the levels of several histone modifications as indicated, in taxane-resistant and sensitive cell lines. Cell extracts were obtained using a mild-modified RIPA buffer. PAC, paclitaxel (20 nM). (C) Heatmap of Pearson correlation scores representing the distribution of the signal intensity of H3K4me1, H3K4me3, H3K27ac and H3K27me3 ChIP-seq in parental and taxane resistant MDA-MB-436 cells showing distinct histone methylation profile in resistant cells relative to parental cells. (D) Heatmap of H3K27me3 ChIP-seq signal intensity in parental and in individual taxane-resistant MDA-MB-436 cells over a 5kb region distributed around the center of common and unique called peaks. Top: venn diagrams showing the number of peaks overlapping and unique for each mark assessed by ChIP-seq. (E) Genomic visualization of H3K27me3 signal over the indicated genomic regions following filtering out the LOCKs regions using CREAM. Top panel represents discrete H3K27me3 genomic regions across parental and the 3 resistant MDA-MB-436 lines; bottom panel represents an example of H3K27me3 LOCKs identified by CREAM across parental and the 3 resistant MDA-MB-436 lines (F) Heatmap of filtered H3K27me3 ChIP-seq signal at discrete genomic regions in parental and in individual resistant cells over a 5 kb region distributed around the center of common and unique called regions upon LOCKS exclusion using CREAM. Top: venn diagrams showing the number of peaks overlapping and unique for each mark assessed by ChIP-seq. (G) Genomic visualization of H3K27me3 signal intensity of parental-specific (left), shared (middle) and resistant-specific LOCKs regions in parental and taxane-resistant MDA-MB-436 cells. Y-axis is normalized signal intensity from ChIP-seq experiment. Assignment of H3K27me3 LOCKs specificity is based on a 2-fold difference in H3K27me3 average signal intensity over the LOCK region between parental and resistant cells. (H) Venn diagram revealing the number of LOCKs regions that are unique to parental cells, unique to taxane-resistant cells or present in both parental and taxane-resistant cells (shared) based on the difference in the average signal intensity of H3K27me3 from ChIP-seq assay over the identified LOCK region using CREAM in resistant cells relative to parental cells (log2FC(Res/P) > 0.05). A cutoff of 2-fold difference in average signal intensity was used to classify the regions. Bottom stacked histogram represents the genomic coverage (in Mb) of each group of LOCKs. (I) Boxplot representing the width distribution (in kb) of the different LOCKs identified by CREAM using a (log2FC(Res/P) > 0.05) for LOCKS classification. (J) Boxplot representing the number of genes expected to be associated to each subtype of LOCKs using randomly generated regions for each parental-specific, shared or resistant-specific LOCKs (Black dot represents the number of genes observed in each subtype of H3K27me3 LOCKs identified in MDA-MB-436 parental and resistant cells. Significance was calculated using permutation test (1000 permutations)

We therefore assessed the genome-wide distribution of H3K4me1, H3K4me3 and H3K27me3 using ChIP-seq assays in all taxane-resistant and parental MDA-MB-436 cells (Figure 4C-D and S4E-F). These were complemented with ChIP-seq assays against H3K27ac (Figure S4G). Pearson correlation between the signal intensity of all populations for each histone modifications shows positive correlations for all active histone modifications (H3K4me1, H3K4me3 and H3K27ac) across taxane-resistant and parental MDA-MB-436 cells (Figure 4C). In agreement, over 83, 63 and 82 % of genomic regions significantly enriched in the taxane-resistant cells for H3K4me1, H3K4me3 or H3K27ac are respectively shared with parental MDA-MB-436 cells while the few resistant- and parental-specific genomic regions are mostly of low ChIP signal intensity (Figure S4E-F-G).

The repressive H3K27me3 modification is anti-correlated with the three active histone modifications (Figure 4C). Furthermore, H3K27me3 ChIP-seq signal is less correlated between taxane-resistant and parental MDA-MB-436 cells, suggesting an important reprogramming of H3K27me3 modification in the taxane-resistant cells relative to parental cells (Figure 4C). Accordingly, between 49 to 53% of the H3K27me3 enriched genomic regions observed in the parental MDA-MB-436 cells are lost in the taxane-resistant cells with only a few regions gaining H3K27me3 signal in the resistant cells (Figure 4D). The reprogramming of H3K27me3 is further characterized by an increase in signal intensity in taxane-resistant cells at genomic regions shared between taxane-resistant and parental MDA-MB-436 cells (Figure 4D). Several of these common H3K27me3 enriched genomic regions cluster into widespread domains reminiscent of LOCKs in taxane-resistant MDA-MB-436 cells(Timp and Feinberg, 2013; Wen et al., 2009). Characterizing the H3K27me3 signals that comprise LOCKs versus single H3K27me3 regions outside of LOCKs using clustering of cis-regulatory regions method (CREAM) (https://www.biorxiv.org/content/early/2017/11/21/222562) in both taxane-resistant and parental MDA-MB-436 cells reveals that over 74% of common H3K27me3 regions, 40% of the parental-specific and 45% of the taxane-resistant-specific H3K27me3 enriched genomic regions fall into LOCKs (Figure 4E-F-G). Importantly, the proportion of H3K27me3 signal comprised within LOCKs rises from an average of 57% of total H3K27me3 genomic regions in parental cells to an average of 68% in taxane-resistant cells, suggestive of taxane-resistant H3K27me3 signal reprogramming towards LOCKs regions (Figure 4F-G). The parental-specific, shared or resistant-specific H3K27me3 discrete genomic regions that are respectively lost, maintained or gained in taxane-resistant cells do not significantly associate with a concomitant change in proximal gene expression (Figure S4H), arguing for a minimal impact of the loss of H3K27me3 signal at discrete genomic regions on gene expression in taxane-resistant MDA-MB-436 cells.

To understand the role of H3K27me3 LOCKs, we classified them using a 2-fold difference cutoff between average H3K27me3 signal in parental and taxane-resistant MDA-MB-436 cells and identified 90 taxane-resistant-specific and 64 parental-specific LOCKs in addition to 659 shared LOCKs (Figure 4H). Taxane-resistant-specific LOCKs are wider than parental-specific or shared LOCKs with an average size of 630 kb in taxane-resistant versus 29 and 21 kb in parental- and shared LOCKs, respectively (Figure 4H). Accordingly, the 90 taxane-resistant-specific LOCKs cover close to 81Mb of the genome in total, which represents about half of the genomic coverage of the 659 shared LOCKs (159Mb), highlighting the widespread nature of H3K27me3 reprogramming into LOCKs in taxane-resistant MDA-MB-436 cells (Figure 4I). Interrogation of the genes comprised within LOCKs reveals that taxane-resistant-specific LOCKs lie in gene-poor regions, a feature common with shared LOCKs but distinct from parental-specific LOCKs (FDR<0.05 by permutation testing using random clusters) (Figure 4J). This demonstrates that the large scale reprogramming of H3K27me3 defines a new epigenetic state characterized by a gain of widespread heterochromatin domains over gene-poor LOCK regions in taxane-resistant MDA-MB-436 cells, concomitant with the loss of a large subset of discrete H3K27me3 genomic regions in taxane-resistant TNBC cells, with no significant effect on gene expression.

### H3K27me3 LOCKs compensate DNA hypomethylation at TE in taxane-resistant TNBC cells

DNA hypomethylation and H3K27me3 LOCKs observed in taxane-resistant cells respectively fall within intergenic and gene-poor genomic regions. Given the well-described repressive nature of DNA methylation and H3K27me3 modification, we asked whether the reprogramming of both epigenetic modifications is orchestrated to maintain a repressive chromatin state over the hypomethylated DNA regions observed in taxane-resistant cells. Focusing on the 41,529 differentially methylated CpGs between taxane-resistant and parental MDA-MB-436 cells (delta-beta-value cutoff +/- 0.3 and P < 0.05), more than 35% lie within H3K27me3 LOCKs (14,628 out of 41,529) (Figure 5A). This represents over 26% of the total number of CpG probes covered by LOCKs regions (14,628/55,842) and is in contrast with the mere 3% (26,901/738,148) of CpG probes located outside of LOCKs (Figure 5A). Furthermore, over 95% (13,949 of 14,628) of the differentially methylated CpG probes located inside H3K27me3 LOCKs are significantly hypomethylated in taxane-resistant cells while probes located outside LOCKs are as likely to be hypo- or hyper-methylated in taxane-resistant MDA-MB-436 cells versus parental cells (54% hypomethylated and 46% hypermethylated) (Figure 5B). The significant hypomethylation associated to H3K27me3 LOCKs in taxane-resistant MDA-MB-436 cells is observed in all parental-specific, resistant-specific and shared LOCKs regions (Figure 5C).

**Figure 5.**
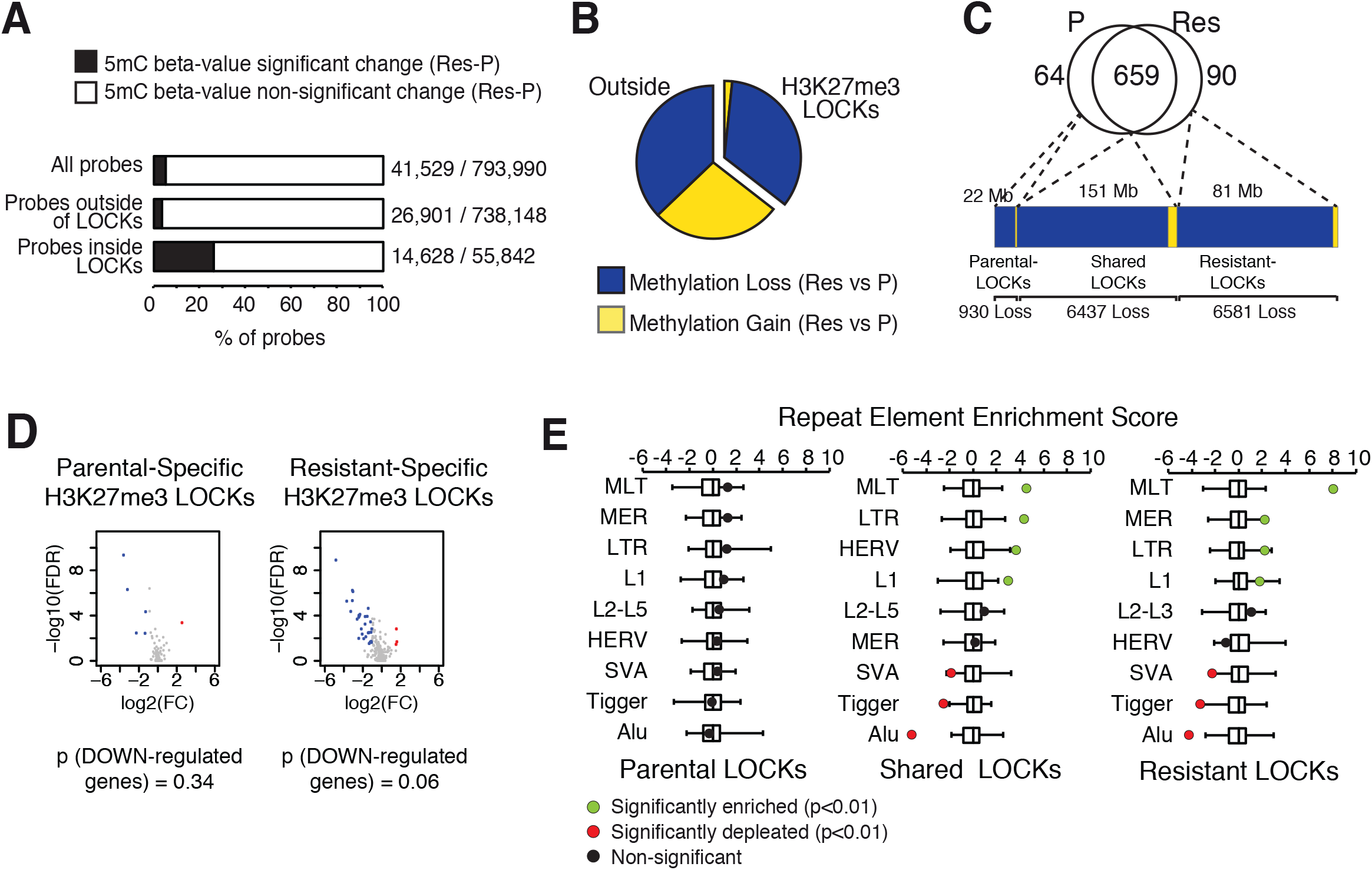
H3K27me3 LOCKs compensate DNA hypomethylation at TE in taxane-resistant TNBC cells. (A) Stacked histogram representing the percentage of probes showing significant alteration of DNA methylation in taxane-resistant cells relative to parental cells for either all probes from the array (top), probes located outside of H3K27me3 LOCKs (middle) or probes located inside the LOCKs regions (bottom). (B) Pie chart representing the 5mC loss (blue) or 5mC gain (yellow) in taxane-resistant cells relative to parental cells for significantly changed 5mC probed located either outside or inside LOCKs regions. (C) Stacked histogram representing the 5mC loss (blue) and 5mC gains (yellow) in each of the parental-specific, resistant-specific or shared LOCKs regions. (D) Volcano plot representing the relative gene expression of genes included in the different H3K27me3 LOCKs (unique to parental or unique to resistant based CREAM calls with a 2-fold fold difference between signal intensity in parental and resistant cells). Blue dots represent LOCKs genes significantly downregulated and red represent genes significantly up-regulated in taxane-resistant cells relative to parental cells. BH-adjusted p-value, FDR<0.05. Significant association between gene expression and H3K27me3-associated genomic regions was calculated using Wilcoxon ranked sum test. (E) Boxplot representing the number of each TE elements expected to be associated to each subtype of LOCKs using randomly generated regions for each parental-specific, shared or resistant-specific LOCKs (100 permutations). Dots represent the number of TE observed in each subtype of H3K27me3 LOCKs identified in MDA-MB-436 parental and resistant cells. Green and red dots respectively show significant enrichment or depletion of TE, which were calculated using 10,000 permutation tests (FDR <0.01).

The preferential reprogramming of H3K27me3 modification at widespread LOCKs is likely providing a compensatory mechanism for maintaining heterochromatin transcriptional repression over these gene-poor DNA-hypomethylated genomic regions (Figure S5A). Accordingly, while a trend towards repression was noted, the change in the expression of genes comprised within H3K27me3 LOCKs was not significant (Figure 5D, p-value downregulated genes = 0.06) between taxane-resistant and parental MDA-MB-436 cells. This suggests that the reprogramming of H3K27me3 to LOCKs enriched for hypomethylated CpGs in taxane-resistant cells, could contribute to maintain transcriptional repression of the few genes comprised in these regions. Given that shared and taxane-resistant LOCKs are significantly prominent at gene-poor regions, we assessed whether they are enriched for other genomic features characteristic of intergenic regions such as TE. Indeed, shared and taxane-resistant-specific H3K27me3 LOCKs are significantly enriched for specific families of TE, including MLT, LINE-1, LTR and MER (p<0.01) while parental-specific LOCKs do not associate with enrichment of any of the transposable element families assessed (Figure 5E). Altogether, this shows that H3K27me3 modification is preferentially reprogrammed into widespread LOCKs over hypomethylated CpGs, providing a compensatory mechanism for maintaining repressive chromatin state over regions enriched for TE.

### Reprogramming of H3K27me3 creates epigenetic vulnerability in taxane-resistant TNBC cells

Expression of endogenous retroviral (ERV) elements originating from TE can induce double-stranded RNA (dsRNA) accumulation, which favors the interferon signaling-mediated activation of the viral mimicry response and blocks cell growth(Liu et al., 2016; Roulois et al., 2015). We assessed whether reprogramming of H3K27me3 serves to maintain the transcriptional repression at TE in absence of DNA methylation in the taxane-resistant MDA-MB-436 cells, thereby preventing ERV expression. We first show that 3 and 5 days exposure to UNC1999, an inhibitor of the EZH2 methyltransferase responsible for H3K27me3 deposition, is equally able to deplete H3K27me3 signal in both parental and taxane-resistant MDA-MB-436 cells (Figure 6A and S6A). Inhibition of H3K27me3 using UNC1999 treatment for 5 days leads to the accumulation of dsRNA in taxane-resistant MDA-MB-436 cells, while no significant dsRNA induction is observed upon UNC1999 treatment in the parental TNBC cells (Figure 6B). Treatment with UNC2400, the control chemical probe for EZH2 inhibition, caused a modest increase in dsRNA at 5-days, specifically in taxane-resistant cells, suggesting hypersensitivity of the taxane-resistant cells to EZH2 inhibition (Figure 6B). In addition, depletion of H3K27me3 following the pharmacological inhibition of EZH2 with UNC1999 induced the expression of specific ERVs from the MER and MLT families in taxane-resistant MDA-MB-436 cells while having no effect on parental cells (Figure 6C).

**Figure 6.**
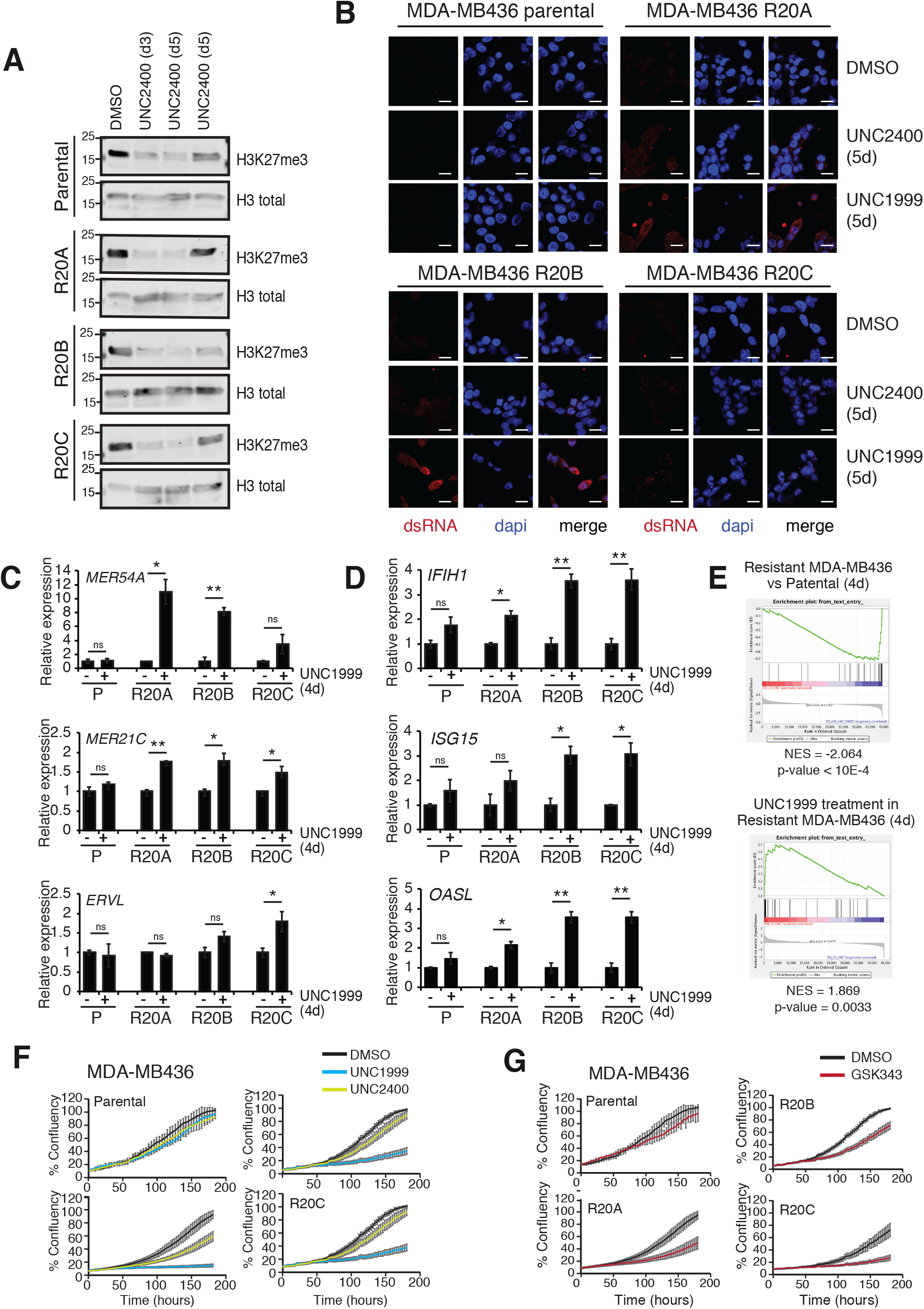
Reprogramming of H3K27me3 at hypomethylated LOCKs regions protects taxane-resistant cells against paclitaxel-induced ds-RNA-mediated interferon response. (A) Western blot showing the levels of H3K27me3 and total H3 in parental and taxane-resistant MDA-MB-436 cells upon 3 and 5 days exposure to the EZH2 inhibitor UNC1999 (3 μM). (B) Immunofluorescence imaging of parental and taxane-resistant MDA-MB-436 cells for detection of dsRNA (red) upon 5-day treatment with EZH2 inhibitor UNC1999 (3 μM) or the control probe UNC2400 (3 μM). Red signal, dsRNA (J2-ab); blue, dapi. (C) RT-QPCR showing the relative expression of the RNA transcripts MER54A, MER21C and ERVL in parental and taxane-resistant MDA-MB-436 cells upon treatment with the EZH2 inhibitor UNC1999 for 96 hours (3 μM). Scale bar represent 20 μm. Statistical analysis Error bars: std deviations. *, p<0.05; **, p<0.01, two-sided t-test. (D) RT-QPCR showing the relative expression of genes comprised in the viral mimicry response in parental and taxane-resistant MDA-MB-436 cells upon 96h treatment with the EZH2 inhibitor UNC1999 (3 μM). Statistical analysis Error bars: std deviations. *, p<0.05; **, p<0.01, paired student t-test. (E) Gene-Set Enrichment Analysis showing the significant association of the interferon response gene signature in the taxane-resistant MDA-MB-436 in presence of paclitaxel relative to parental cells (middle panel) or upon 4 days treatment with UNC1999 in taxane-resistant cells (3 μM). (F) Time-dependant proliferation assay (IncucyteZoom) of taxane-resistant and sensitive cells MDA-MB-436 upon treatment with the EZH2 inhibitors UNC1999 (3 μM) or the control probe UNC2400 (3 μM). Resistant cells were grown in constant presence of paclitaxel (20 nM). Error bars: std deviations. (G) Time-dependant proliferation assay (IncucyteZoom) of taxane-resistant and sensitive cells MDA-MB-436 upon treatment with the EZH2 inhibitors GSK343 (3 μM). Resistant cells were grown in constant presence of paclitaxel (20 nM). Error bars: std deviations.

We asked whether accumulation of dsRNA upon alleviation of the H3K27me3 modification in the taxane-resistant MDA-MB-436 cells can induce of the expression of genes regulating the interferon-response pathway. Accordingly, induction of interferon-response gene expression is observed specifically in taxane-resistant cells upon treatment with UNC1999 (Figure 6D). Comparison of gene expression profiles between parental and taxane-resistant TNBC cells reveal that upon development of taxane-resistance, the TNBC cells adopt a gene expression profile that is negatively associated with the interferon-response gene signature (P < 1e-04, NES=-2.064) (Figure 6E). Treatment of the resistant cells with UNC1999, through the accumulation of dsRNA, induces in a gene expression profile that positively associates with the activation of the interferon response signature (P = 0.0033, NES=1.869) (Figure 6E).

Finally, we assessed whether alleviating the H3K27me3-mediated repression at LOCKs using EZH2 inhibition creates an epigenetic vulnerability in taxane-resistant MDA-MB-436 cells. While inhibition of EZH2-mediated H3K27me3 modification using UNC1999 or GSK343 has no effect on the growth and survival of parental MDA-MB-436 or Hs 578T cells, it significantly antagonized the growth and survival of all taxane-resistant MDA-MB-436 and Hs 578T cells (Figure 6F-G and S6B-D), thereby creating an epigenetic vulnerability. Accordingly, dose-response cell growth assay shows that all taxane-resistant TNBC cells gained sensitivity to lower concentrations of EZH2 inhibitors compared to parental cells (Figure S6E). Together, these results demonstrate that the taxane-resistance in TNBC cells dictates an epigenetic profile that warrants the maintenance of a repressive heterochromatin-state over vulnerable LOCKs regions in order to prevent the induction of ERVs expression and concomitant accumulation of dsRNA and activation of interferon-response.

## DISCUSSION

Diverse metabolic adaptations support the development of drug-resistant cancer cells under the stressful environment that stems from therapy(Bhola et al., 2016; Komurov et al., 2012; Lopes-Rodrigues et al., 2017). Here, we show that a systematic impairment in methionine metabolism across all taxane-resistant TNBC cells, giving rise to increased methionine dependency in the drug-resistant models. The observed increase in SAM-consuming metabolites including choline, 1-MNA, carnitine and activation of transulfuration in taxane-resistant TNBC cells parallels previous observations in drug-resistant cancer cells. In particular, transsulfuration and glutathione metabolism ensures proper metabolic detoxification capacity in cancer cells exposed to drugs(Deblois et al., 2016; Ju et al., 2015; Kerr et al., 2016). In addition, methylation of phophatidylethanolamine (PE) is a major consumer of SAM for the synthesis of phosphatidylcholine(Ye et al., 2017) and aberrant choline phospholipid metabolism was associated with tamoxifen resistance in breast cancer cell lines(Kim et al., 2017). Likewise, carnitine metabolism favours cancer cells survival upon conditions of metabolic stress such as glucose deprivation or hypoxia(Barger et al., 2013; Zaugg et al., 2011). Therefore, the accumulation of SAM-consuming metabolites may ensure metabolic conditions favoring the survival of taxane-resistant TNBC cells in toxic drug environment.

An important consequence of the altered methionine metabolism observed in taxane-resistant TNBC cells is the global decrease in DNA methylation compensated by the acquisition of H3K27me3 LOCKs, specifically occurring over TEs. This parallels previous studies showing that altered one-carbon metabolism resulting from genetic alterations in metabolic enzymes or from methionine-restricted diets associate with global changes in LINE-1 methylation(Friso et al., 2002; Kloypan et al., 2015; Kottakis et al., 2016; Llanos et al., 2015). The fact that global decreases in DNA methylation observed in taxane-resistant TNBC cells is not paralleled with a global decrease in H3K27me3 levels agrees with the higher Michaelis constant (Km) for SAM reported for DNA methyltransferases (DNMTs) compared to EZH2, suggesting that DNMTs activity is more susceptible to changes in SAM levels(Bacolla et al., 2001; Horiuchi et al., 2013; Meier, 2013). However, the sensitivity of different methyltransferases to changes in SAM availability also depends on cell type and developmental stage(Yun et al., 2012).

Given that DNA hypomethylation is an early event in carcinogenesis, and accentuates in advanced disease(Timp et al., 2014), the spreading of the H3K27me3 LOCKs formation over hypomethylated DNA could be essential for cancer cells progression and survival upon drug exposure. Our observations parallel reports showing compensation of DNA hypomethylation by the accumulation of H3K27me3 in mouse embryonic stem cells to ensure genome stability(Walter et al., 2016) and of a panel of repressive histone methylation (H3K27me3, H3K9me3 and H4K20me3) in various cancer types, to maintain transcriptional repression of TEs(Hon et al., 2012; Statham et al., 2012; Varley et al., 2013). Uncompensated DNA hypomethylation can reactivate the expression of TEs, including ERVs. giving rise to accumulation of dsRNAs. This activates the interferon-induced viral mimicry response, which inhibits cell growth(Beauregard et al., 2008; Liu et al., 2016; Roulois et al., 2015). In agreement, repression of TEs is essential for immune escape in leukemic stem cells(Colombo et al., 2017) and for the survival of drug-tolerant lung cancer cell subpopulations(Guler et al., 2017). The compensation of DNA hypomethylation over TEs by H3K27me3 LOCKs in taxane-resistant TNBC cells highlights the importance of transcriptionally repressing TEs for the taxane-resistance in our TNBC models. The requirement for evading the viral mimicry response to acquire resistance to taxane is further supported by the downregulation of the interferon response genes in taxane-resistant TNBC cells. Accordingly, alleviating the compensation of DNA hypomethylation by H3K27me3 using EZH2 inhibitors leads to accumulation of dsRNA and activation of the interferon-response to block cell growth.

The various mechanisms proposed to underlay the development of resistance to standard-of-care treatment in cancer are diverse, yet most of them involve the engagement of survival pathways that bypass or compensate for the detrimental effects induced by the drugs. Together, this study supports a role for metabolic adaptations in inducing epigenetic reprogramming allowing for the development of taxane-resistant TNBC cells by ensuring the maintained repression of TEs. Importantly, this epigenetic reprogramming generates an epigenetic vulnerability, that can be targeted with chemical inhibitors of EZH2 to re-sensitize taxane-resistant TNBC cells.

## MATERIAL & METHODS

### Cell culture, reagents and antibodies

For all experiments, MDA-MB-436 and Hs 578-t were maintained in high-glucose DMEM (Gibco) supplemented with 10% foetal bovine serum (FBS, Gibco) and Penicilin/Streptomycin (100 U/mL Penicilium and 100 μg/mL. Streptomycin) routinely tested for mycoplasma. Parental cells were obtained from ATCC. Taxol-resistant cells were derived as previously described upon long-term exposure to increasing concentrations of paclitaxel at a starting concentration 0.05 nM with increments (dose and time adjusted to the growth and survival of adapting cells), until the cytotoxic concentration of 10 to 20 nM was reached with the resistant cells growing at a similar rate than the parental cells (>6 months). All cell lines were routinely tested for mycoplasma Resistant cells were continuously cultured in the presence of 10 or 20 nM paclitaxel (Sigma-Aldrich, T4702), a cytotoxic concentration for the parental TNBC cells. The same volume of DMSO (Cederlane, DMS666.100) was used as a control in the maintenance of the parental TNBC cells. For L-methionine depletion and deprivations, taxol-parental and resistant cells were transferred in custom DMEM supplemented with dialyzed FBS deprived of glucose, glycine, glutamine, serine and methionine (Wisent, 319-026-CL) supplemented with L-glucose (Sigma, G7021), 30 mg/L glycine (Sigma, G8790), 42 mg/L serine and 200 mM glutamine (Sigma, S4311). For long-term L-methionine deprivation, cells were maintained in custom DMEM supplemented with dialyzed FBS deprived of glucose, glycine, glutamine, serine and methionine (Wisent, 319-026-CL) supplemented with L-glucose (Sigma, G7021), 30 mg/L glycine (Sigma, G8790), 42 mg/L serine and 200 mM glutamine (Sigma, S4311) and supplemented with 3 mg/L L-methionine (10% of L-methionine levels in regular DMEM) for more than 4 weeks. For methionine, homocysteine, SAM add-back experiments, methionine-deprived DMEM was supplemented with 30 mg/L or 3 mg/L L-methionine (Sigma, M8439), 250 μM SAM (Sigma, A4377), or 0.4 mM L-homocysteine (Sigma, 69453). For acute paclitaxel treatment, parental cells were transferred to DMEM containing 20 nM paclitaxel for the indicated times. For transfection of poly(i:C), poly(i:C) was diluted in LyoVec (Invitrogen, lyec-2) transfection reagent 15 minutes prior to transfection and added to the cells for 24h prior harvesting for RNA-preparation or immunostaining. ChIP assays were performed using an anti-H3K4me1 rabbit polyclonal antibody (Abcam Ab8895), H3K4me4 mouse monoclonal antibody (Millipore 05-1339), H3K27ac rabbit polyclonal antibody (Abcam Ab4729) and H3K27me3 rabbit polyclonal antibody (Diagenode C15410069). Western blots on cell extracts were performed using the following antibodies: anti-beta-actin mouse monoclonal antibody (SC47778), anti-H3K4me1 rabbit polyclonal antibody (Abcam Ab8895), anti-H3K4me4 mouse monoclonal antibody (Millipore 05-1339), anti-H3K27ac rabbit polyclonal antibody (Abcam Ab4729), anti-H3K27me3 rabbit polyclonal antibody (Diagenode C15410069), anti-H3 rabbit polyclonal antibody (Abcam, AB1791), anti-H3K27me1 (EMD-Millipore 07-448) and anti-H3K36me3 (Abcam) rabbit polyclonal antibody.

### Metabolomic profiling and analysis

Metabolomics analysis was performed with liquid chromatography (LC) coupled to Q Exactive Plus high resolution mass spectrometer (HRMS). The HPLC (Ultimate 3000 UHPLC)was coupled to QE-MS (Thermo Scientific) for metabolite separation and detection. An Xbridge amide column (100 × 2.1 mm i.d., 3.5 μm; Waters) was employed for compound separation at room temperature. Mobile phase A was water with 5 mM ammonium acetate (pH 6.9), and mobile phase B was acetonitrile. The gradient was linear as follows: 0 min, 85% B; 1.5 min, 85% B; 5.5 min, 35% B; 10 min, 35% B; 10.5 min, 35% B; 10.6 min, 10% B; 12.5 min, 10% B; 13.5 min, 85% B; and 20 min, 85% B. The follow rate was 0.15 ml/min from 0 to 5.5 min, 0.17 ml/min from 6.9 to 10.5 min, 0.3 ml/min from 10.6 to 17.9 min, and 0.15 ml/min from 18 to 20 min. For SAM analysis, mobile phase A was replaced with water containing 10 mM ammonium acetate (pH8.5). HRMS acquisition and raw data analysis method was described previously (Analytical chemistry 86 (4), 2175-2184). For intracellular metabolite analysis, 5e^5^ cells were first incubated in 6 well plate in 3 mL medium for 96 hours. Cells were briefly washed with 1 ml ice cold saline (0.9 % NaCl in water) twice before metabolite extraction. Metabolites were extracted into 80 % methanol, dried in a vacuum concentrator, and stored in −80 °C freezer until further analysis. Medium metabolites were extracted into 80 % methanol and was directly subjected to LC-MS analysis. Intracellular metabolites were reconstituted into sample solvent (water:methanol:acetonitrile, 2:1:1, v/v) and the volume of sample solvent was proportional to cell number, and 3 μl was injected for metabolomics analysis. The LC-MS peak intensity (integrated peak area) of each metabolite was used to calculate the relative abundance ratio. Peak intensity values obtained from HRLC-MS were quantile normalized and differential enrichment of metabolites was calculated between taxane-resistant TNBC cells and parental cells using one-way ANOVA. Metabolic set enrichment analysis for differentially expressed metabolites (MSEA) and pathway enrichment for integrated genes expression and metabolites levels were assessed with Metaboanalyst server.

### Global DNA methylation quantification, DNA methylation array and analysis

Quantification of 5-mC DNA was assayed using the Global DNA Methylation Assay Line-1 ELISA system (Active Motif) as specified by the commercant. Bisulphite conversion of the DNA for methylation profiling was performed using the EZ DNA Methylation kit (Zymo Research) on 500 ng genomic DNA from 4 replicates of each parental and taxane-resistant cell lines. Conversion efficiency was quantitatively assessed by quantitative PCR (qPCR). The Illumina Infinium MethylationEPIC BeadChips were processed as per manufacturer’s recommendations. The R package ChAMP v2.6.4 was used to process and analyze the data for differentially methylation probes and regions(Butcher and Beck, 2015; Morris et al., 2014). For analysis purposes, probes differentially methylated between parental and taxane-resistant MDA-MB-436 cells with p-value <0.05 and delta-beta value < −0.3 or > 0.3 were considered significantly changed. Data have been deposited at GSE111541.

### Protein extraction, analysis and quantification of H3K27me3

Proteins were extracted from 2-3M cells using specified extraction buffers including RIPA lysis buffer: 25 mM Tris•HCl, pH 7.6, 150 mM NaCl, 1% NP-40, 1% sodium deoxycholate, 0.1% sodium dodecyl sulfate (SDS). MILD modified RIPA lysis buffer: 50 mM Tris HCl, pH 8, 150 mM NaCl, 1 mM EDTA, 1% NP-40, 0.25% deoxycholate or acid extraction buffer (0.4M HCl and neutralized with sodium divasic phosphate. Two to 10 μg of proteins were loaded on precast polyacrylamide gel (AnyKD, Biorad) and analyzed by western blotting. Quantification of H3K27me3 in TNBC cells was achieved using the Histone H3 methylated Lys27 Elisa system (Active Motif). Histones were isolated using an acid extraction for Elisa.

### ChIP-sequencing

For ChIP-sequencing analyses, chromatin was prepared from parental and taxol-resistant MDA-MB-436 and Hs 578T cells grown at 70-80% confluence. For each immunoprecipitation, 3-5M cells were crosslinked with 1% formaldehyde for 10 minutes at room temperature. Crosslink was stopped using cold PBS washes and cells were pelleted and lysed and extracted chromatin was sonicated as previously described(McGuirk et al., 2013). ChIP was performed using 2 μg of antibody directed towards each Histone modifications reported. The ChIP primers are listed in Supplementary Table 1. For ChIP-sequencing, Sonicated chip DNA samples were quantified using qubit reagent (Life Technologies); one nanogram of DNA per sample was used following ThruPLEX DNA-seq kit following manufacturer’s protocol. Amplification of libraries were monitored in real time for optimal number of cycles. Final libraries were size selected to 260-350 bp using Pippin HT (Sage Science). ChIP libraries were size validated using Agilent Bioanalyzer and concentration validated by qPCR (Kapa Biosystems/Roche). All libraries were normalized to 10 nM and pooled together, denatured with 0.2 N NaOH and diluted to a final concentration of 10 pM. Pooled libraries were loaded onto cBot (Illumina) for cluster generation. Clustered flow cell was then sequenced Single read 50 cycles V3 using Illumina Hiseq2000 to achieve ∼30 million reads per sample. The sequences generated from ChIP-seq were analysed with the following specifications. Reads were filtered for quality (phred33 score >=30) and length (n>=32) trimming using FASTX Toolkit 0.0.13.1. Filtered reads were aligned to the reference assembly hg19 using bwa v.0.5.9. Only uniquely mapped reads were retained for further analysis. Redundant reads were removed using SAMtools v0.1.18. Number of reads were downsampled to the lowest number of reads of the samples to be compared together. Peaks were called using MACS2.0 with default values (mfold[10,30], p-value 1e-04). WIG files were generated using MACS2.0 with default parameters. Bed files contain peak calls on merged reads using corresponding inputs as control. Score represent peak intensity. Pearson correlation of signal intensity was obtained and Peak annotation, tag directory, bed file generation, merging and was performed with the Homer package v3.1 (http://biowhat.ucsd.edu/homer/ngs/index.html). Tag heatmaps were obtained with Java TreeView(Saldanha, 2004). Data have been deposited at insert GEO archive

### CREAM analysis

H3K27me3 LOCKs were identified using CREAM considering cut-off of 0.1 for window-size in both parental and taxane-resistant MDA-MB436 cells using the H3K27me3 peak calls from MACs2.0. The clusters were further annotated as parental-specific, resistant-specific, or shared between parental and taxane-resistant MDA-MB436 cells considering 2 fold-change difference between H3K27me3 average signal intensity within LOCKs in parental and resistant samples.

### RNA extraction, RNA-sequencing, analysis and gene set enrichment

For expression analysis, parental and taxane-resistant TNBC cells were harvested at 80% confluency and RNA extraction was performed with the RNeasy Mini Kit (Qiagen) and reverse transcribed using SensiFAST cDNA synthesis kit (Bioline) and analysed by Q-RT-PCR with SensiFAST SYBR-green NO-ROX kit (Bioline). For gene expression profiling from RNA-sequencing, library preparations were made from RNA samples that were quantified by qubit (Life Technologies) and were quality by Agilent Bioananlyzer. All samples had RIN score >8 were library prepared using TruSeq Stranded mRNA kit (Illumina). Two hundred nanograms total RNA from the 30 RNA samples were purified for polyA tail containing mRNA molecules using poly-T oligo attached magnetic beads, following purification the RNA was fragmented. The cleaved RNA fragments were copied into first strand cDNA using reverse transcriptase and random primers. This is followed by second strand cDNA synthesis using RNase H and DNA Polymerase I. A single “A” based were added and adapter ligated followed by purification and enrichment with PCR to create cDNA libraries. Alternatively, for UNC1999 treatment experiment, RNA samples were quantified by qubit (Life Technologies) and quality by Agilent Bioananlyzer. Two hundred nanograms Total RNA from 42 samples were library prepared using TruSeq Stranded Total RNA kit (Illumina). RNA samples were ribosomal RNA depleted using Ribo-zero Gold rRNA beads, following purification the RNA was fragmented. The cleaved RNA fragments were copied into first strand cDNA using reverse transcriptase and random primers. This is followed by second strand cDNA synthesis using RNase H and DNA Polymerase I. A single “A” based were added and adapter ligated followed by purification and enrichment with PCR to create cDNA libraries. For RNA-sequencing, final cDNA libraries were size validated using Agilent Bioanalyzer and concentration validated by qPCR (Kapa Biosystems/Roche). All libraries were normalized to 10 nM and pooled together, denatured with 0.2 N NaOH and diluted to a final concentration of 8 pM. Pooled libraries were loaded onto cBot (Illumina) for cluster generation. Clustered flow cell was then sequenced Paired-end 100 cycles V3 using Illumina Hiseq2000 to achieve ∼30 million reads per sample. Alternatively for UNC treated samples, final cDNA libraries were size validated using Agilent Bioanalyzer and concentration validated by qPCR (Kapa Biosystems/Roche). All libraries were normalized to 10 nM and pooled together, denatured with 0.2 N NaOH and diluted to a final concentration of 1.4 pM. 1.3 ml of 1.4 pM pooled libraries were loaded onto an Illumina NextSeq cartridge for cluster generation and sequencing on an Illumina Nextseq500 instrument (Illumina) using Paired-end 75bp protocol to achieve ∼ 40 million reads per sample. Reads were aligned to hg19 using tophat (2.0.8) with default parameters, transcripts were merged using cufflinks (2.1.1) and differential expression computed for all relevant contrasts with default parameters. Differentially expressed genes were used as input for determining gene set enrichment analysis (GSEA) using the GSEA using canonical pathways or hallmarks of cancer pathways. Primer sequences for RT-PCR for gene expression are listed in Supplementary Table 2. Data have been deposited at GSE111920 (UNC1999 treatment) and (steady-state gene expression).insert GEO archive.

### Association of ChIP-seq and cluster regions with gene expression

Genes in 100 kb proximity of individual peaks or LOCKs are considered as genes associated to those regions. Expression of these genes obtained from RNA-sequencing were then compared between parental and taxane-resistant samples using Wilcoxon ranked sum test. The resulting p-values were FDR corrected for all the hypothesis (number of genes) tested for each set of elements (set of peaks or LOCKs identified for a given ChIP-Seq profile).

### Repeat element enrichment score

To identify the enrichment of each family of repeat elements, we first obtained sets of randomly selected chromosome regions with the same distribution of than the LOCKs within each chromosome. The actual number of repeats identified in LOCKs for either parental-specific, shared or resistant-specific H3K27me3 LOCKs were compared with the number of repeats obtained in the 10,000 randomly selected chromosome regions. FDR was calculated as one minus the number of times the random regions contained more (positive enrichment) or less (negative enrichment) repeat elements than the observed H3K27me3 LOCKs and divided by the total number of random set of regions (10,000).

### Cell proliferation assays

Cells were maintained as described above, seeded in 24-well plates and treated with the indicated epigenetic probes or cytotoxic drugs or supplemented with or depleted from the indicated metabolites. Cell viability was assessed using crystal violet as previously described. Alternatively, cells were seeded in 96-well plates transferred to IncuCyte® ZOOM Live cell analysis system (Essen BioScience) and proliferation was monitored using IncuCyte® ZOOM phase-contrast quantification software. All results shown represent the average of at least 3 independent replicate experiments each carried out with at least 3 technical replicates.

### Immunofluorescence imaging

Parental and taxol-resistant TNBC cells were seeded in 6 well plates and treated with UNC1999 (3 μM), UNC2400 (3 μM) or DMSO on the following day for 4 days. Cells were then counted and seeded into 24-well plates at a density of 1×10^5^ per well and treated again with UNC1999 (3 μM), UNC2400 (3 μM) or DMSO for another 24h. Cells were fixed in methanol, and stained overnight with J2 scion antibody for dsRNA staining (Mouse J2 antibody Product No: 10010200 Scicons 1:500) or dapi for nuclear staining (Hoescht, H1399 Thermo Fisher Scientific, 1:2000) and incubated in secondary antibody (Cell Signaling 4410S anti-mouse IgG alexa 647 1:1000) or anti-Rabbit secondary antibody (Cell signaling, alexa 488 1:1000) imaged in using a 40X objective. Total and dsRNA-positive cells in a minimum of 4 independent fields (at least 1000 total cells per condition) were quantified using ImageJ. The experiment was performed three times and statistical significance was determined using Student’s t-test.

## Acknowledgements

We thank Ken Kron, Aislin Treloar, Christopher Arlidge and Sarina Cameron for assistance in the maintenance of the various cell lines. The authors wish to thank the PMH sequencing core facility for sequencing of ChIP and RNA and for methylEPIC array hybridization. This study was conducted with the support of the Terry Fox Research Institute, Canadian Cancer Research Society and the Ontario Institute for Cancer Research through funding provided by the Government of Ontario. This work was also supported by the Canadian Institute for Health Research (CIHR: Funding Reference Number 136963 to M.L. and 363288 to B.H.K.), the Princess Margaret Cancer Foundation (M.L.) and the Gattuso-Slaight Personalized Cancer Medicine Fund at Princess Margaret Cancer Centre (B.H.K.). We acknowledge the Princess Margaret Bioinformatics group for providing the infrastructure assisting us with analysis presented here. G.D. is a recipient of fellowships from the CIHR and of the Fonds de Recherche en Santé du Québec (FRSQ) doctoral research award. M.L. holds an Investigator Award from the Ontario Institute for Cancer Research, a CIHR New Investigator Award and the Bernard and Francine Dorval Award for Excellence from the Canadian Cancer Society. P.G. is a recipient of a CIHR fellowship. S.A.M.T was supported by Connaught International Scholarships for Doctoral Students. The SGC is a registered charity (number 1097737) that receives funds from AbbVie, Bayer Pharma AG, Boehringer Ingelheim, Canada Foundation for Innovation, Eshelman Institute for Innovation, Genome Canada through Ontario Genomics Institute [OGI-055], Innovative Medicines Initiative (EU/EFPIA) [ULTRA-DD grant no. 115766], Janssen, Merck KGaA, Darmstadt, Germany, MSD, Novartis Pharma AG, Ontario Ministry of Research, Innovation and Science (MRIS), Pfizer, São Paulo Research Foundation-FAPESP, Takeda, and Wellcome.

