## Supplementary Materials for "Metabolic adaptations underlie epigenetic vulnerabilities in chemoresistant breast cancer"

#### INVENTORY OF SUPPLEMENTARY INFORMATION

**Supplementary Figures and legends to Supplementary Figures 1-6**

**Supplementary Tables 1-2**

### Figure S1

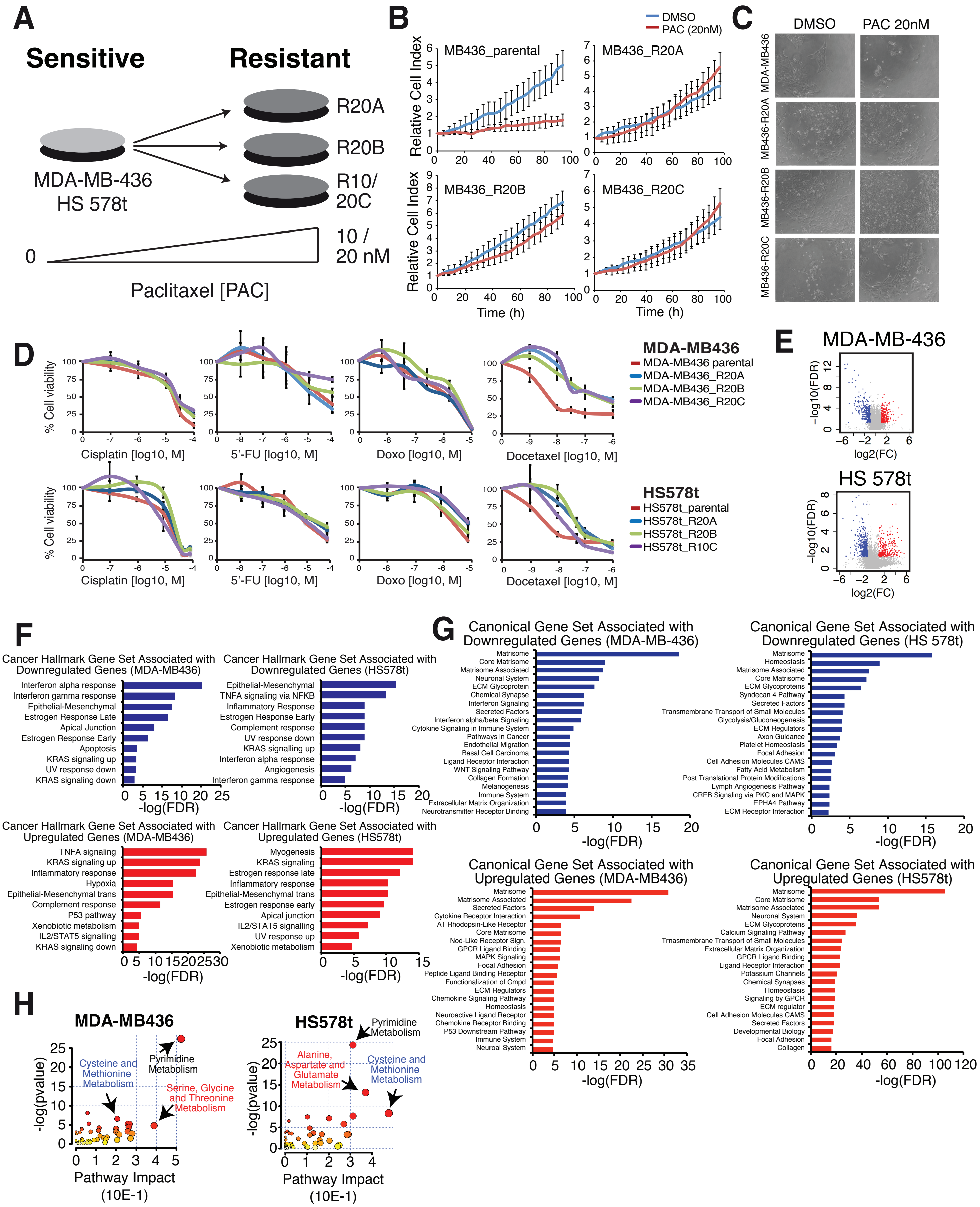

A
Sensitive
Resistant
MDA-MB-436
HS 578t
R20A
R20B
R10/
20C
0
10 /
20 nM
Paclitaxel [PAC]
B
DMSO
PAC (20nM)
MB436_parental
MB436_R20A
MB436_R20B
MB436_R20C
Relative Cell Index
Time (h)
C
DMSO
PAC 20nM
MDA-MB436
MB436-R20A
MB436-R20B
MB436-R20C
D
% Cell viability
Cisplatin [log10, M]
5'-FU [log10, M]
Doxo [log10, M]
Docetaxel [log10, M]
MDA-MB436
MDA-MB436 parental
MDA-MB436_R20A
MDA-MB436_R20B
MDA-MB436_R20C
HS578t
HS578t_parental
HS578t_R20A
HS578t_R20B
HS578t_R10C
E
MDA-MB-436
HS 578t
−log10(FDR)
log2(FC)
F
Cancer Hallmark Gene Set Associated with Downregulated Genes (MDA-MB436)
Interferon alpha response
Interferon gamma response
Epithelial-Mesenchymal
Estrogen Response Late
Apical Junction
Estrogen Response Early
Apoptosis
KRAS signaling up
UV response down
KRAS signaling down
Cancer Hallmark Gene Set Associated with Downregulated Genes (HS578t)
Epithelial-Mesenchymal
TNFA signaling via NFKB
Inflammatory Response
Estrogen Response Early
Complement response
UV response down
KRAS signalling up
Interferon alpha response
Angiogenesis
Interferon gamma response
Cancer Hallmark Gene Set Associated with Upregulated Genes (MDA-MB436)
TNFA signaling
KRAS signaling up
Inflammatory response
Hypoxia
Epithelial-Mesenchymal trans
Complement response
P53 pathway
Xenobiotic metabolism
IL2/STAT5 signalling
KRAS signaling down
Cancer Hallmark Gene Set Associated with Upregulated Genes (HS578t)
Myogenesis
KRAS signaling
Estrogen response late
Inflammatory response
Epithelial-Mesenchymal trans
Estrogen response early
Apical junction
IL2/STAT5 signalling
UV response up
Xenobiotic metabolism
-log(FDR)
G
Canonical Gene Set Associated with Downregulated Genes (MDA-MB-436)
Matrisome
Core Matrisome
Matrisome Associated
Neuronal System
ECM Glycoprotein
Chemical Synapse
Interferon Signaling
Secreted Factors
Interferon alpha/beta Signaling
Cytokine Signaling in Immune System
Pathways in Cancer
Endothelial Migration
Basal Cell Carcinoma
Ligand Receptor Interaction
WNT Signaling Pathway
Collagen Formation
Melanogenesis
Immune System
Extracellular Matrix Organization
Neurotransmitter Receptor Binding
Canonical Gene Set Associated with Downregulated Genes (HS 578t)
Matrisome
Homeostasis
Matrisome Associated
Core Matrisome
ECM Glycoproteins
Syndecan 4 Pathway
Secreted Factors
Transmembrane Transport of Small Molecules
Glycolysis/Gluconeogenesis
ECM Regulators
Axon Guidance
Platelet Homeostasis
Focal Adhesion
Cell Adhesion Molecules CAMS
Fatty Acid Metabolism
Post Translational Protein Modifications
Lymph Angiogenesis Pathway
CREB Signaling via PKC and MAPK
EPHA4 Pathway
ECM Receptor Interaction
Canonical Gene Set Associated with Upregulated Genes (MDA-MB436)
Matrisome
Matrisome Associated
Secreted Factors
Cytokine Receptor Interaction
A1 Rhodopsin-Like Receptor
Core Matrisome
Nod-Like Receptor Sign.
GPCR Ligand Binding
MAPK Signaling
Focal Adhesion
Peptide Ligand Binding Receptor
Functionalization of Cmpd
ECM Regulators
Chemokine Signaling Pathway
Homeostasis
Neuroactive Ligand Receptor
Chemokine Receptor Binding
P53 Downstream Pathway
Immune System
Neuroal System
Canonical Gene Set Associated with Upregulated Genes (HS578t)
Matrisome
Core Matrisome
Matrisome Associated
Neuronal System
ECM Glycoproteins
Calcium Signaling Pathway
Trnasmembrane Transport of Small Molecules
Extracellular Matrix Organization
GPCR Ligand Binding
Ligand Receptor Interaction
Potassium Channels
Chemical Synapses
Homeostasis
Signaling by GPCR
ECM regulator
Cell Adhesion Molecules CAMS
Secreted Factors
Developmental Biology
Focal Adhesion
Collagen
-log(FDR)
H
MDA-MB436
Cysteine and Methionine Metabolism
Pyrimidine Metabolism
Serine, Glycine and Threonine Metabolism
HS578t
Pyrimidine Metabolism
Alanine, Aspartate and Glutamate Metabolism
Cysteine and Methionine Metabolism
-log(pvalue)
Pathway Impact (10E-1)

### Figure S2

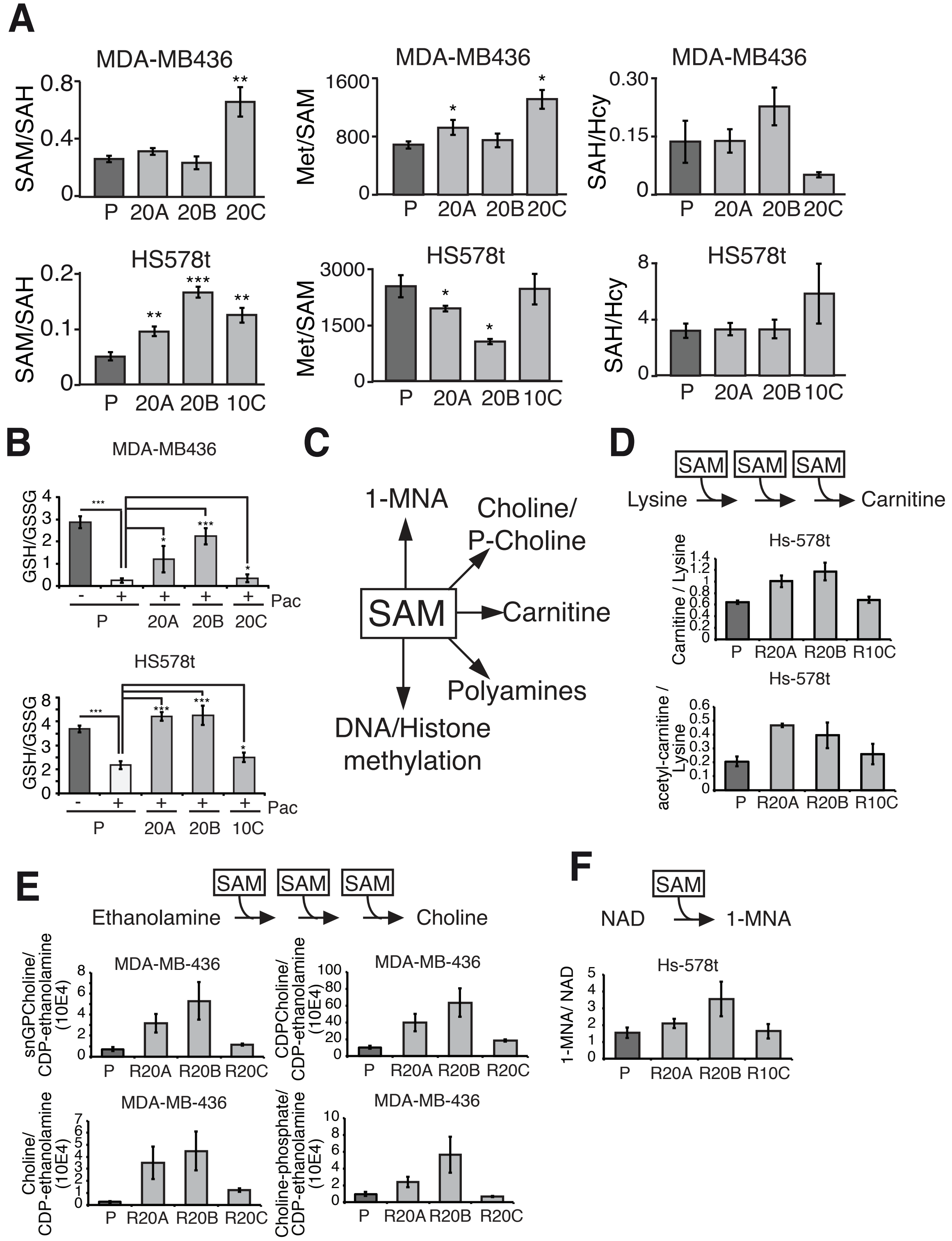
A
MDA-MB436
SAM/SAH
Met/SAM
SAH/Hcy
P
20A
20B
20C
HS578t
10C
B
MDA-MB436
GSH/GSSG
Pac
HS578t
C
1-MNA
Choline/
P-Choline
SAM
Carnitine
Polyamines
DNA/Histone
methylation
D
Lysine
Carnitine
Hs-578t
Carnitine / Lysine
acetyl-carnitine /
Lysine
R20A
R20B
R10C
E
Ethanolamine
Choline
MDA-MB-436
snGPCholine/
CDP-ethanolamine
(10E4)
CDPCholine/
CDP-ethanolamine
Choline/
Choline-phosphate/
R20C
F
NAD
1-MNA
1-MNA/ NAD

### Figure S3

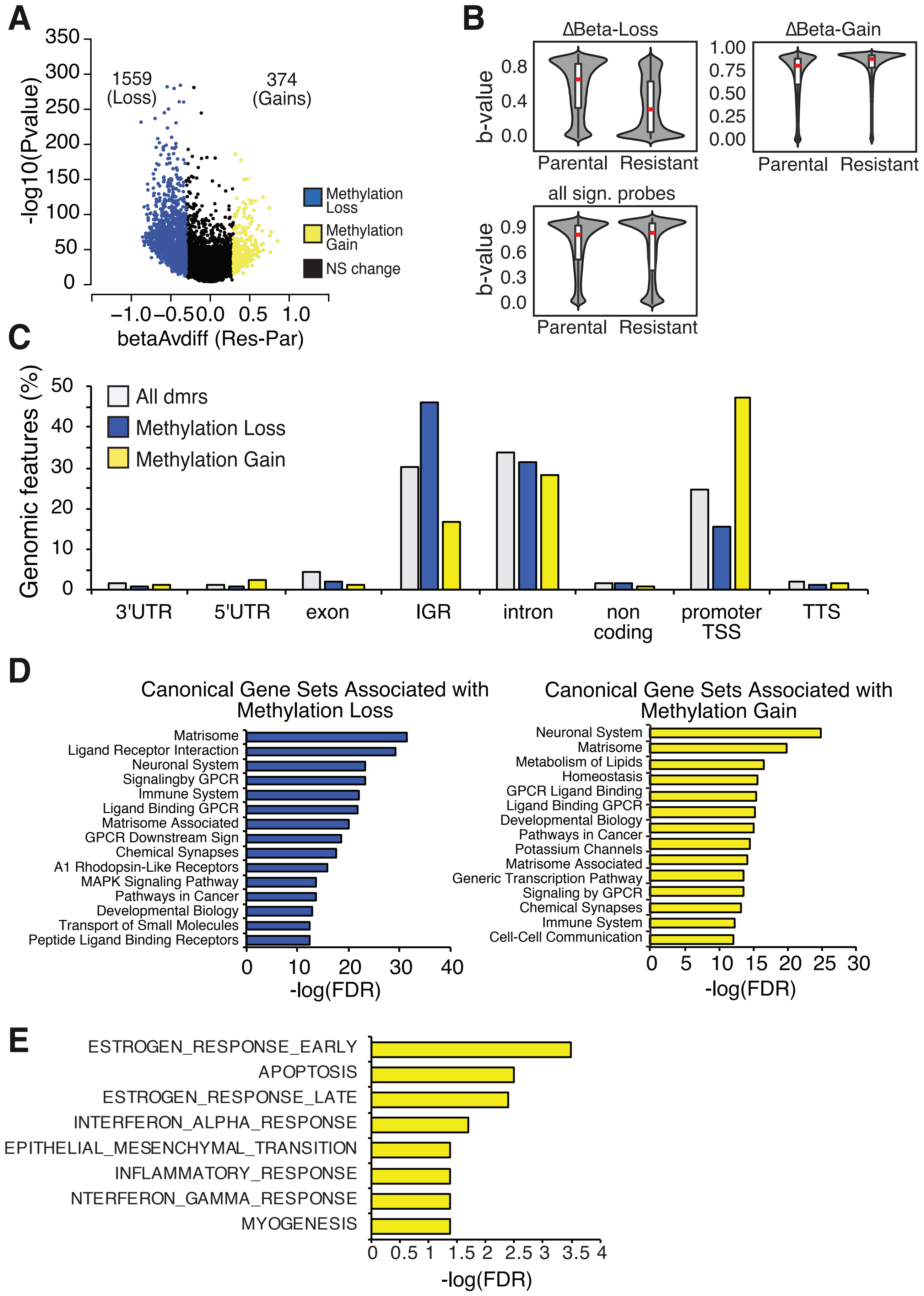

A
-log10(Pvalue)
350
300
250
200
150
100
50
0
1559
(Loss)
374
(Gains)
Methylation Loss
Methylation Gain
NS change
−1.0 −0.5 0.0 0.5 1.0
betaAvdiff (Res-Par)
B
ΔBeta-Loss
ΔBeta-Gain
b-value
0.8
0.4
0.0
1.00
0.75
0.50
0.25
0.00
Parental
Resistant
all sign. probes
0.9
0.6
0.3
0.0
C
Genomic features (%)
All dmrs
Methylation Loss
Methylation Gain
50
40
30
20
10
0
3'UTR
5'UTR
exon
IGR
intron
non coding
promoter TSS
TTS
D
Canonical Gene Sets Associated with Methylation Loss
Matrisome
Ligand Receptor Interaction
Neuronal System
Signalingby GPCR
Immune System
Ligand Binding GPCR
Matrisome Associated
GPCR Downstream Sign
Chemical Synapses
A1 Rhodopsin-Like Receptors
MAPK Signaling Pathway
Pathways in Cancer
Developmental Biology
Transport of Small Molecules
Peptide Ligand Binding Receptors
0 10 20 30 40
-log(FDR)
Canonical Gene Sets Associated with Methylation Gain
Neuronal System
Matrisome
Metabolism of Lipids
Homeostasis
GPCR Ligand Binding
Ligand Binding GPCR
Developmental Biology
Pathways in Cancer
Potassium Channels
Matrisome Associated
Generic Transcription Pathway
Signaling by GPCR
Chemical Synapses
Immune System
Cell-Cell Communication
0 5 10 15 20 25 30
-log(FDR)
E
ESTROGEN_RESPONSE_EARLY
APOPTOSIS
ESTROGEN_RESPONSE_LATE
INTERFERON_ALPHA_RESPONSE
EPITHELIAL_MESENCHYMAL_TRANSITION
INFLAMMATORY_RESPONSE
NTERFERON_GAMMA_RESPONSE
MYOGENESIS
0 0.5 1 1.5 2 2.5 3 3.5 4
-log(FDR)

### Figure S4

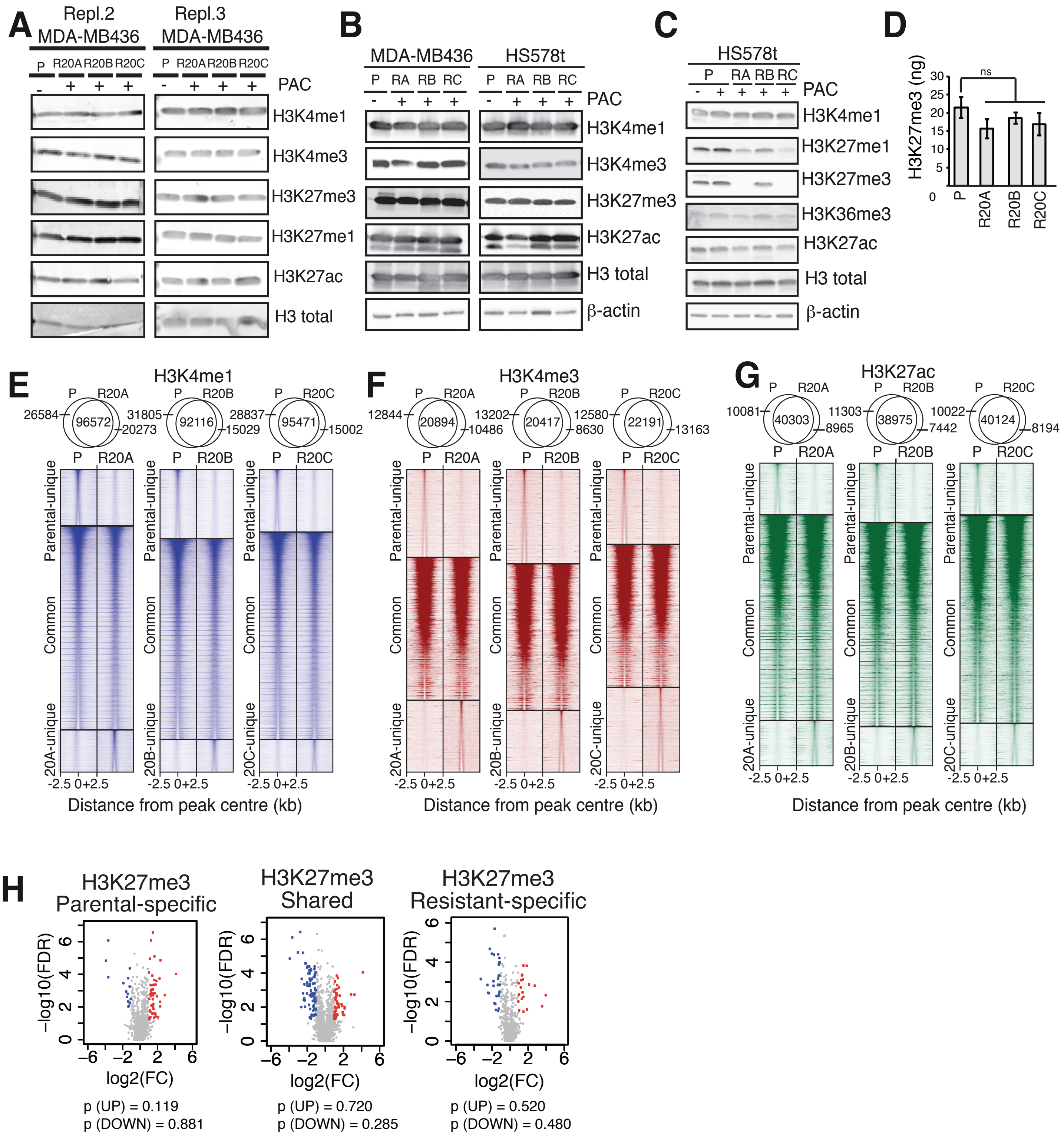
A
Repl.2 MDA-MB436
Repl.3 MDA-MB436
P R20A R20B R20C
PAC
H3K4me1
H3K4me3
H3K27me3
H3K27me1
H3K27ac
H3 total
B
MDA-MB436
HS578t
P RA RB RC
PAC
H3K4me1
H3K4me3
H3K27me3
H3K27ac
H3 total
β-actin
C
HS578t
P RA RB RC
PAC
H3K4me1
H3K27me1
H3K27me3
H3K36me3
H3K27ac
H3 total
β-actin
D
H3K27me3 (ng)
ns
P R20A R20B R20C
E
H3K4me1
26584 96572 20273
31805 92116 15029
28837 95471 15002
Parental-unique
Common
20A-unique
20B-unique
20C-unique
-2.5 0+2.5
Distance from peak centre (kb)
F
H3K4me3
12844 20894 10486
13202 20417 8630
12580 22191 13163
G
H3K27ac
10081 40303 8965
11303 38975 7442
10022 40124 8194
H
H3K27me3 Parental-specific
H3K27me3 Shared
H3K27me3 Resistant-specific
−log10(FDR)
log2(FC)
p (UP) = 0.119
p (DOWN) = 0.881
p (UP) = 0.720
p (DOWN) = 0.285
p (UP) = 0.520
p (DOWN) = 0.480

### Figure S5

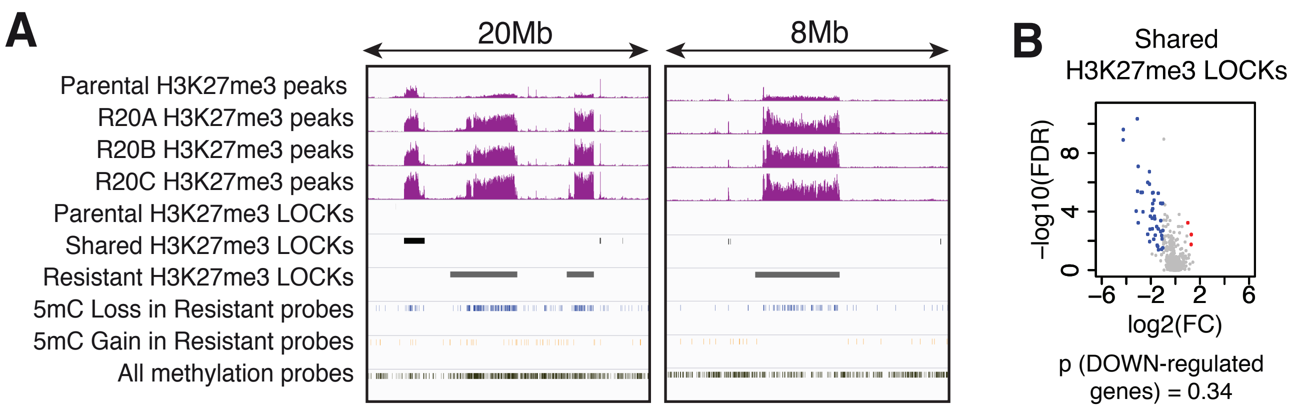

A
20Mb
8Mb
Parental H3K27me3 peaks
R20A H3K27me3 peaks
R20B H3K27me3 peaks
R20C H3K27me3 peaks
Parental H3K27me3 LOCKs
Shared H3K27me3 LOCKs
Resistant H3K27me3 LOCKs
5mC Loss in Resistant probes
5mC Gain in Resistant probes
All methylation probes
B
Shared
H3K27me3 LOCKs
−log10(FDR)
0
4
8
−6
−2
2
6
log2(FC)
p (DOWN-regulated
genes) = 0.34

### Figure S6

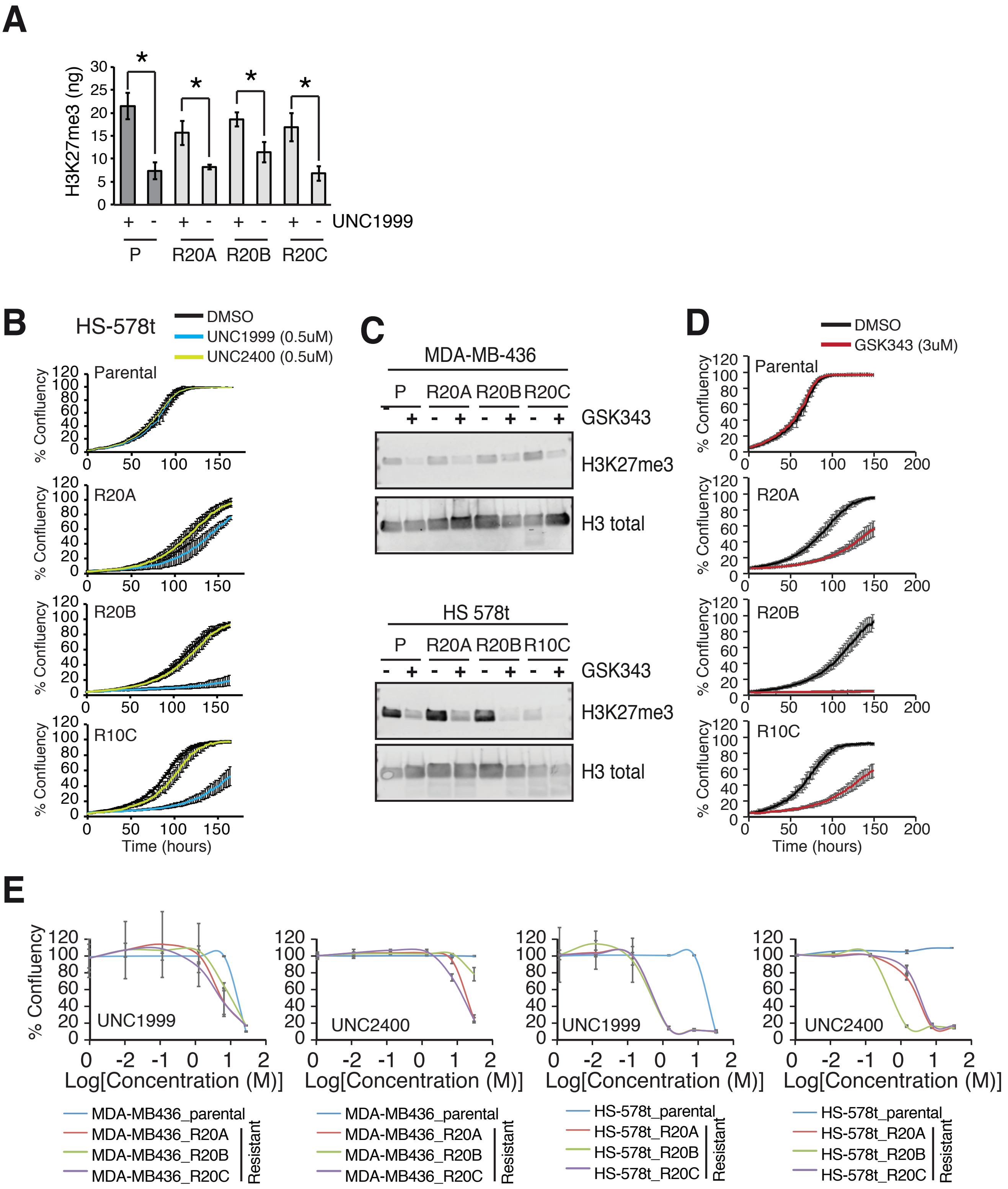

A
H3K27me3 (ng)
UNC1999
P
R20A
R20B
R20C
B
HS-578t
DMSO
UNC1999 (0.5uM)
UNC2400 (0.5uM)
Parental
R20A
R20B
R10C
% Confluency
Time (hours)
C
MDA-MB-436
HS 578t
P
R20A
R20B
R20C
R10C
GSK343
H3K27me3
H3 total
D
DMSO
GSK343 (3uM)
Parental
R20A
R20B
R10C
% Confluency
Time (hours)
E
% Confluency
UNC1999
UNC2400
Log[Concentration (M)]
MDA-MB436_parental
MDA-MB436_R20A
MDA-MB436_R20B
MDA-MB436_R20C
HS-578t_parental
HS-578t_R20A
HS-578t_R20B
HS-578t_R20C
Resistant

#### SUPPLEMENTARY FIGURES

**Figure S1 |**

(A) Taxane-resistance was developed in TNBC cells upon long-term exposure to increasing concentration of paclitaxel ranging from 0 to 20 nM. (B) Time-dependant proliferation curves (IncucyteZoom) upon paclitaxel treatment (20 nM, red curve) in parental and taxane-resistant MDA-MB-436 cells. Error bars: stdev (C) Picture (IncucyteZoom) of the parental and resistant MDA-MB-436 cells upon paclitaxel treatment (20 nM). (D) Dose-dependence concentration curves showing the % cell viability response of the parental and taxane-resistant cells upon exposure to 72h of other standard-of-care therapeutic drugs at indicated concentrations. Error bars: stdev (E) Volcano plot representing the relative gene expression of for taxane-resistant cells relative to parental cells in MDA-MB-436 and Hs 578T cells. Blue dots represent genes significantly downregulated; red represent genes significantly up-regulated in taxane-resistant cells relative to parental cells. Gray dots are genes not significantly changed. BH-adjusted p-value, FDR<0.05. (F) Bar graph representing gene ontology enrichment of significant Cancer Hallmark pathways for genes either significantly upregulated (dark grey) or significantly downregulated (light grey) in resistant cells relative to parental cells. FDR<0.05. (G) Bar graph representing gene ontology enrichment of significant Canonical pathways for genes either significantly upregulated (dark grey) or significantly downregulated (light grey) in resistant cells relative to parental cells. FDR<0.05. (H) Pathway enrichment of differentially expressed metabolites between parental and taxane-resistant TNBC cells combining overrepresentation analysis (Hypergeometric Test) and Pathway topology analysis (Relative-betweenness

Centrality). Yellow, non-significant; Red, significant enrichment. Size of dots correspond to higher impact pathways.

**Figure S2 |**

(A) Relative ratio of relative abundance in steady-state levels of successive metabolites of the methionine cycle in parental and taxane-resistant TNBC cells. Error bars: std. Statistical significance is calculated using 2-sided unpaired t-test; * denotes $p<0.05$; ** denotes $p<0.01$. (B) Relative conversion ratio of steady-state levels of GSH/GSSG taxane-resistant parental and resistant TNBC cells. Error bars: std. Statistical significance is calculated using 2-sided unpaired t-test; * denotes $p<0.05$; ** denotes $p<0.01$. (C) Schematic representation of various fates of SAM towards SAM-consuming metabolic pathways. (D) Conversion rations of subsequent metabolites in the generation of carnitine from lysine in Hs 578T parental and resistant cells. Error bars: std, statistical significance calculated by two-sided unpaired t-test, * denotes $p<0.05$; ** denotes $p<0.01$. (E) Conversion rations of subsequent metabolites in the conversion generation of choline-derivatives from ethanolamine derivatives in MDA-MB-436 parental and resistant cells. Error bars: std, statistical significance calculated by two-sided unpaired t-test, * denotes $p<0.05$; ** denotes $p<0.01$. (F) Conversion rations of subsequent metabolites in the conversion generation of 1-MNA from nicotinamide in MDA-MB-436 parental and resistant cells. Error bars: std, statistical significance calculated by two-sided unpaired t-test, * denotes $p<0.05$; ** denotes $p<0.01$.

**Figure S3 |**

(A) Volcano plot showing significant differentially methylated regions identified by Probe Lasso in taxane-resistant relative to parental MDA-MB-436 cells (delta-beta-value (Resv-P)); ($p<0.05$). Each dot represents a differentially methylated regions (dmrs), blue dots represent significantly hypomethylated CpGs with delta-beta value < -0.3 (Res-P), yellow dots represent significantly hypomethylated CpGs with delta-beta value > +0.3 (Res-P), black dots represent non-significantly (NS) altered CpGs with delta-beta values between -0.3 and +0.3. (B) Violin plots representing the distribution of delta-beta values (Res-P) for probes in parental and taxane-resistant MDA-MB-436 cells for either all CpGs (bottom panel), for hypermethylated CpGs (top right panel) or for hypomethylated CpGs (top left panel)  (C) Percentage of distribution of genomic features associated to all dmrs (white), to the dmrs showing a significant loss of methylation in the resistant cells (blue) or to the dmrs showing a significant gain of methylation in resistant cells relative to parental cells. (D) Canonical gene-set enrichment associated to upregulated genes with promoter dmr that show 5mC loss (blue) or with downregulated genes that have promoter dmrs that show 5mC gain in the resistant cells relative to parental cells. (E) Hallmark of cancer gene-set enrichment associated to downregulated genes that have promoter dmrs that show 5mC gain in the resistant cells relative to parental cells. $FDR<0.05$.

**Figure S4 |**

(A) Western blot depicting the levels of several histone modifications as indicated, in taxane-resistant and sensitive cell lines. Cell extract was obtained using ChIP lysis buffer (complete cell extract) used to perform ChIP-seq experiment. PAC, paclitaxel (20 nM). (B) Western blot depicting the levels of assessed histone modifications as indicated, in taxane-resistant and sensitive Hs 578T lines. Cell extracts were obtained using a RIPA-ChIP buffer. PAC, paclitaxel (20nM). (C) Western blot depicting the levels of assessed histone modifications as indicated, in taxane-resistant and sensitive Hs 578T lines. Cell extracts were obtained using a mild-modified RIPA buffer. PAC, paclitaxel (20 nM). (D) Quantification of H3K27me3 levels in parental (P) and taxane-resistant (R) MDA-MB-436 cells H3K27me1/3 quantification ELISA system using recombinant Histone 3 (H3) as standards. Heatmap of H3K4me1 (E), H3K4me3 (F) and H3K27ac (G) ChIP-seq signal intensity in parental and in individual resistant cells over a 5 kb region distributed around the center of common and unique called peaks. Top: venn diagrams showing the number of peaks overlapping and unique for each mark assessed by ChIP-seq. Venn diagram representing the lost, shared or gained H3K27me3 regions in parental (P) or taxane-resistant (Res) MDA-MB-436 cells. Volcano plot representing the relative gene expression of genes included in the different H3K27me3 peaks (unique to parental, unique to resistant or shared based peak caller). Blue dots represent genes significantly downregulated and red represent genes significantly up-regulated in taxane-resistant cells relative to parental cells. BH-adjusted p-value, $FDR<0.05$. Significant association between gene expression and H3K27me3-associated genomic regions was calculated using Wilcoxon ranked sum test.

**Figure S5 |**

(A) Genomic visualization of H3K27me3 signal intensity of various LOCKs regions in parental and taxane-resistant MDA-MB-436 cells. Y-axis is normalized signal intensity from ChIP-seq experiment. Below are tracks showing H3K27me3 LOCKs identified as resistant-, parental-specific or shared and tracks representing the significantly hypomethylated (blue) or hypermethylated (yellow) CpGs over the genomic region. The track for all probes present on the array is shown at the bottom (black). (B) Volcano plot representing the relative gene expression of genes included in the shared H3K27me3 LOCKs (shared based CREAM calls with a 2-fold fold change in H3K27me3 signal intensity between parental and resistant cells). Blue dots represent LOCKs genes significantly downregulated and red represent genes significantly up-regulated in taxane-resistant cells relative to parental cells. BH-adjusted p-value, FDR<0.05. Significant association between gene expression and H3K27me3-associated genomic regions was calculated using Wilcoxon ranked sum test.

**Figure S6|**

(A) Quantification of H3K27me3 levels in parental (P) and taxane-resistant (R) MDA-MB-436 cells upon 72h exposure to the EZH2 inhibitor UNC1999 (3 μM) using the H3K27me1/3 quantification ELISA system using recombinant Histone 3 (H3) as standards. (B) Time-dependant proliferation assay (IncucyteZoom) of taxane-resistant and sensitive cells Hs

578T upon treatment with 3 μM EZH2 inhibitors UNC1999 (0.5 μM). UNC2400 (0.5 μM) is used as the control probe. The resistant cells were grown in constant presence of paclitaxel (20 nM). Error bars: std deviations. (C) Western blot depicting the levels of H3K27me3 as indicated, in taxane-resistant and sensitive cell lines upon treatment with GSK343 (3 μM). Cell extract was obtained using acid extraction buffer. PAC, paclitaxel (20 nM). (D) Time-dependant proliferation assay (IncucyteZoom) of taxane-resistant and sensitive cells Hs 578T upon treatment with 3 μM GSK343 inhibitors UNC1999 (0.5 μM). The resistant cells were grown in constant presence of paclitaxel (20 nM). Error bars: std deviations. (E) Concentration-dependant proliferation assay (IncucyteZoom) of taxane-resistant and sensitive MDA-MB-436 and Hs 578T cells upon treatment with EZH2 inhibitors (3 μM) at 4 days post-treatment. The resistant cells were grown in constant presence of paclitaxel (20 nM). Error bars: std deviations.

**Supplementary Table1**
**ChIP primers – List of the primers used for histone modification ChIP**

| | |
|---|---|
| H3K27me3-pos-F | GTG AGT CTC TGA GCA GCA CC |
| H3K27me3-pos-R | GGA TGG TGG CTG GAG CTT G |
| H3K27me3-pos2-F | GCC ACC AGA TCG TAT CTC CTG |
| H3K27me3-pos2-R | GCC GAC TGC GGT GAA GTC |
| H3K4me3-pos1-R | GAAGAGGCGGACCCAGCGGT |
| H3K4me3-pos1-R | TTCCTGCCGGTCATCTCGCTT |
| H3K4me3-neg1-R | TGGGTGGTGTCATCTGGTAA |
| H3K4me3-meg1-R | GGATGGAATGGATCAGATGG |
| H3K4me3-neg2-R | GGAGAGGGTGGACAGATTGA |
| H3K4me3-neg2-R | TCAGCCTTGCTGCTACAGTG |
| H3K4me3-pos2-R | GCCAATCTCAGTCCCTTCCC |
| H3K4me3-pos2-R | TAGTAGCCGGGCCCTACTTT |

**Supplementary Table1**
**ChIP primers – List of the primers used for validation of histone modification ChIP**

| | |
|---|---|
| H3K27me3-pos-F | GTG AGT CTC TGA GCA GCA CC |
| H3K27me3-pos-R | GGA TGG TGG CTG GAG CTT G |
| H3K27me3-pos2-F | GCC ACC AGA TCG TAT CTC CTG |
| H3K27me3-pos2-R | GCC GAC TGC GGT GAA GTC |
| H3K4me3-pos1-R | GAAGAGGCGGACCCAGCGGT |
| H3K4me3-pos1-R | TTCCTGCCGGTCATCTCGCTT |
| H3K4me3-neg1-R | TGGGTGGTGTCATCTGGTAA |
| H3K4me3-meg1-R | GGATGGAATGGATCAGATGG |
| H3K4me3-neg2-R | GGAGAGGGTGGACAGATTGA |
| H3K4me3-neg2-R | TCAGCCTTGCTGCTACAGTG |
| H3K4me3-pos2-R | GCCAATCTCAGTCCCTTCCC |
| H3K4me3-pos2-R | TAGTAGCCGGGCCCTACTTT |
